# A Hypothalamic Pathway for Energy Conservation

**DOI:** 10.64898/2026.10.06.756421

**Authors:** Amelia M. Douglass, Nicole Lynch, Matias Andina, Oscar D. Ramirez-Plascencia, Hakan Kucukdereli, David Melville, Luis H. A. Costa, Yuelin Yang, Chenxi Qiu, Sathyajit S. Bandaru, Alexander S. Banks, Sinisa Hrvatin, Clifford B. Saper, Bradford B. Lowell, Natalia L.S. Machado

**Affiliations:** Division of Endocrinology, Metabolism, and Diabetes, Beth Israel-Deaconess Medical Center and Harvard Medical School, Boston 02215 USA; Department of Neurology, Beth Israel Deaconess Medical Center and Harvard Medical School, Boston 02215 USA; Department of Brain and Cognitive Sciences, Massachusetts Institute of Technology, Cambridge 02139 USA; Whitehead Institute for Biomedical Research, Cambridge 02142 USA; Department of Biology, Massachusetts Institute of Technology, Cambridge 02139 USA; Institute of Science and Technology Austria (ISTA), Am Campus 1, 3400 Klosterneuburg, Austria

## Abstract

Many animals utilize a survival strategy of hypothermia, inactivity, and a dramatic decrease in their energy metabolism when faced with a shortfall in energy supplies. Mice, for example, show daily torpor that may last several hours to bridge the lack of energy availability in a cold environment. While our recent studies identified a population of neurons in the median preoptic nucleus (MnPO) necessary for fasting-induced torpor regulation, the specific brain circuits that trigger energy conservation in response to food restriction are unknown.

We show here that the energy state of the animal is conveyed to preoptic neurons by inputs from the arcuate nucleus (ARC), a region known to be indispensable for driving metabolic and behavioral adaptations to the fasted state. We first found that the activity of ARC neurons expressing Agouti-Related Peptide (ARC^AgRP^) is required for the generation of fasting-induced torpor. Then, we showed that torpor-active ARC^AgRP^ neurons innervate the preoptic area of the hypothalamus (POA), including the MnPO. Using optogenetic methods, we also found that activating the ARC^AgRP^→MnPO pathway induces key features of torpor: decreased body temperature, bradycardia, reduction of energy expenditure, and suppression of brown adipose tissue thermogenesis. Remarkably, we also show that inhibition of the ARC^AgRP^→MnPO brain circuit entirely prevents fasting-induced torpor. Furthermore, we identified that both NPY and GABA are neuromodulators used by ARC^AgRP^ neurons to enable hypothermic responses during starvation. Our studies provide a detailed view of a hypothalamic pathway enabling energy conservation, a survival mechanism in conditions of low energy availability.

## Introduction

Thermoregulation plays a vital role in the survival of animals, maintaining core body temperature (Tb) in a narrow range over the course of the day. This sophisticated and energetically costly system evolved in mammalian species, allowing thermoregulatory and metabolic adaptations to seasons of food abundance or periods of starvation^1–3^. Some animals use torpor as a mechanism to suppress energy expenditure (EE) and prolong survival during periods when food supplies are limited and external temperatures are too low to sustain Tb around 37 °C^1^. Humans also show adaptive responses to calorically restrictive diets, including reduced Tb, cold intolerance^4^, and decreased basal metabolic rate (BMR)^5^. However, the basis for this adaptive modulation of the thermoregulatory and metabolic control is still enigmatic.

Mice fed an *ad libitum* (AL) diet ordinarily attempt to maintain a tight control of Tb in a cool environment (the thermoneutral point for these animals is about 29 °C ambient temperature during the light phase and 32 °C at the dark phase^6^, below which additional heat production is required to defend normothermic levels of Tb). However, if kept at ambient temperature (Ta) of about 22 °C and treated with systemic injection of the hunger hormone ghrelin during their less active (sleep or light photoperiod) phase, they develop a hypothermic response^7^. Mice kept in cool environments (such as 22° C) and fasted overnight go into torpor, and this regulated hypothermic and hypometabolic state can be exacerbated by systemic ghrelin treatment^8^. Ghrelin-deficient mice also show disruptions in their ability to produce torpor bouts^9^, indicating the critical role of hunger signaling in inducing the reduction of Tb observed during fasting periods^10^.

Previous studies also suggest that non-specific ablation of neurons in the arcuate nucleus of the hypothalamus (ARC), a region that (1) contains neurons that express ghrelin receptors and (2) provides signals in response to nutritional deficit^9,11–13^, prevents torpor in fasted mice^8,9^. However, the identity of torpor-active ARC neurons and the brain circuits that trigger a reduction in Tb and energy conservation in response to food deprivation (FD) remain unknown.

Neurons located in the preoptic region of the hypothalamus (POA) that express the genes *Adyacp1*, *Qrfp*, and *Esr1*^14–16^ regulate fasting-induced torpor. Silencing these neurons can delay the onset of torpor and alter the structure or degree of the hypothermic response that is typical of torpor bouts. However, identifying a specific genetic marker of glutamatergic (*Slc17a6*, Vglut2) neurons required for the daily torpor phenomenon^14^ has been challenging. Nevertheless, we have recently reported that ablating neurons in the median preoptic nucleus that express the prostaglandin E2 receptor EP3 (MnPO^EP3R^ neurons) is sufficient to prevent mice from generating fasting-induced torpor bouts. Activating MnPO^EP3R^ neurons also causes a dramatic drop in Tb of mice with a reduction of EE and heart rate (HR) to about 20-30% of normal^17^. During this time, mice do not eat, but also do not lose weight. When they recover euthermia, they spontaneously resume activity and regular behavioral responses. The relatively small cluster of MnPO^EP3R^ neurons expresses *Slc17a6, Adcyap1*, *Qrfp,* and *Esr1* genes^18,19^, suggesting that the neurons that uniquely express this cluster of genes are the key population that causes torpor responses^17^. However, for the MnPO^EP3R^ neurons to cause adaptive responses that include hypothermia and hypometabolic states in response to FD, they must receive signals concerning the animals’ metabolic state^10^.

We hypothesized that the energy state of animals is conveyed to thermoregulatory MnPO^EP3R^ neurons by inputs from the ARC. As ARC neurons that express Agouti-Related Peptide (ARC^AgRP^) are activated in fasted animals, we further hypothesized that fasting-activated ARC^AgRP^ neurons drive energy conservation during periods of food unavailability or in response to nutritional deficits by modulating the activity of the MnPO^EP3R^ neurons, which are required for torpor generation.

### ARC^AgRP^ neurons are active at torpor entry and are necessary for torpor generation

Disruption of signaling via the orexigenic neuropeptide Y (NPY) has been shown to reduce the depth and length of daily torpor in mice^8^. As it is well established that NPY is a neuropeptide released by ARC^AgRP^ neurons to control many adaptations to the fasted state, including reducing EE^12,13,20–23^, we hypothesized that AgRP/NPY-expressing ARC neurons are a critical population involved in the generation of fasting-induced torpor. To test this, we first measured changes in intracellular Ca^2+^ levels in ARC^AgRP^ neurons, as a proxy for neural activity, using fiber photometry *in vivo.* To do so, we injected AAV-GCaMP6s into the ARC of *Agrp*-IRES-cre mice. Then, we monitored the changes in calcium signaling during torpor (Fig. 1A). Torpor can be induced in C57BL6 mice at 22 °C Ta by the removal of food access during an overnight period^24^. Initial fasting-related food seeking behavior is marked by increased locomotor activity, a transient Tb elevation, and endogenous glucose production^25^. However, as this effect wanes, mice show reproducible torpor events (periods where Tb usually reaches < 33 °C for at least 1 hour) near the onset of the following light phase^17,24^. Our recording of both ARC^AgRP^ neuron activity and Tb during fasting revealed that ARC^AgRP^ neuronal activity and Tb are inversely related (Ext data Fig 1). During torpor, the calcium signaling recorded from ARC^AgRP^ neurons increased (Fig. 1B, C). Elevated ARC^AgRP^ neuron activity during torpor (Fig 1D-E) suggests that these neurons may be involved with the generation of torpor bouts.

**Fig. 1.**
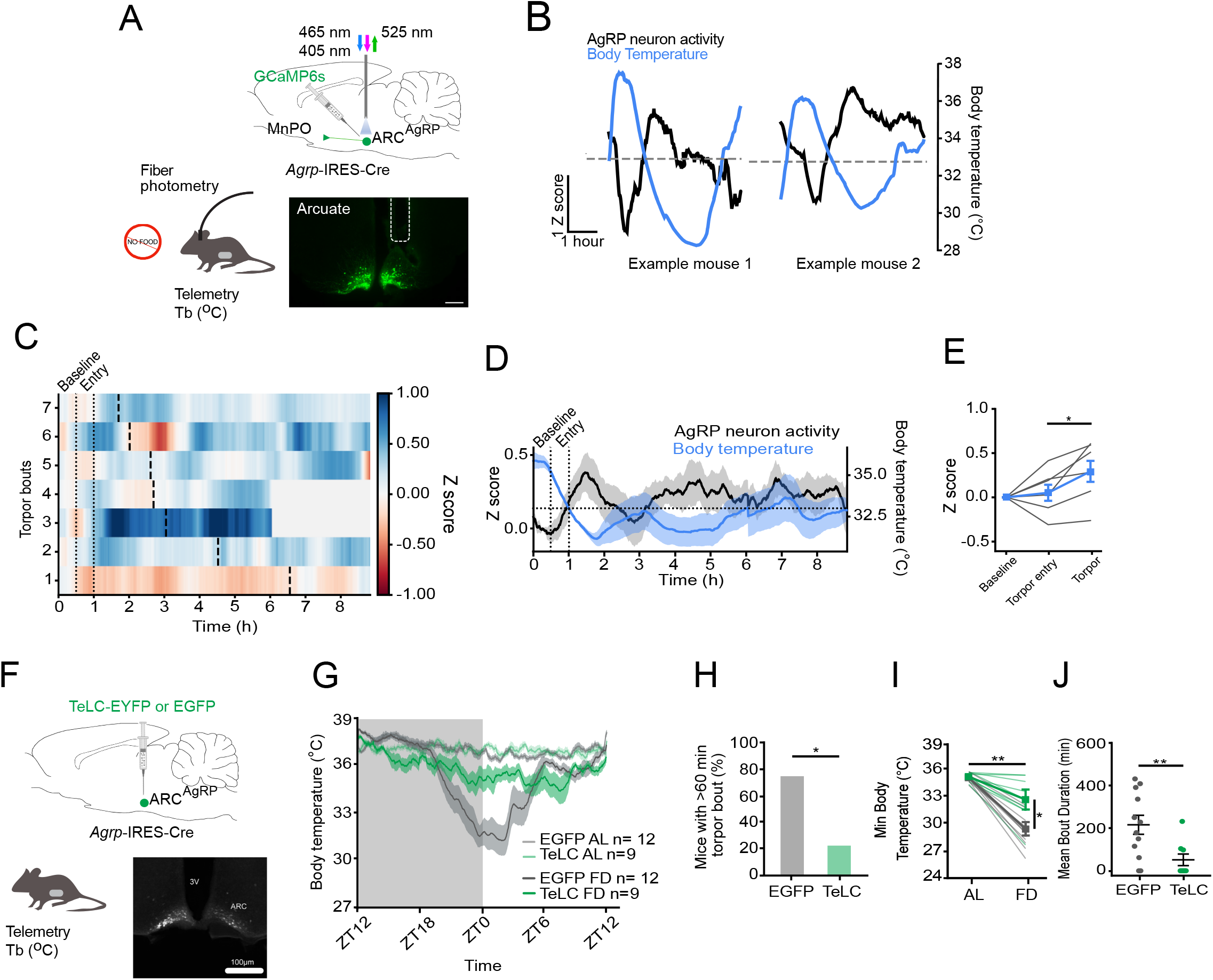
ARC^AgRP^ neurons are required for fasting-induced torpor. (A) Schematic of *in vivo* fiber photometry recordings from ARC^AgRP^ neurons (top) concurrent with biotelemetry recordings of Tb (lower left) and a representative image of GCaMP in ARC^AgRP^ neurons and placement of the optic fiber in the arcuate (lower right). Scale bar = 200 μm (B) Representative traces of ARC^AgRP^ neuron activity and Tb during topor bouts. Dotted horizontal lines: topor, defined as < 33 °C. (C) Heatmap of ARC^AgRP^ activity during all recorded torpor bouts (n=7 bouts from 3 mice), arranged by bout length. Thick vertical black lines = end of the bout. Grey shaded periods denote bouts where the recording had ended. (D) Mean AgRP neuron activity and Tb during torpor bouts. Dotted horizontal lines: topor, defined as < 33 °C. (E) Mean AgRP neuron activity prior to the entry to torpor, during the entry and during torpor bouts (n=7 torpor bouts from 3 mice). Gray lines: individual mice. One-way repeated measures ANOVA with Tukey’s multiple comparisons test, \**p*=0.0204 (F) Schematic of experimental design for ARC^AgRP^ neuron inhibition via TeLC delivery to the arcuate nucleus (top), concurrent with biotelemetry recordings of body temperature (lower left) and a representative image of viral expression in AgRP neurons (lower right). Scale bar = 250 μm (G) Mean body temperature recorded from ARC^AgRP-TeLC^ (TeLC, green) or ARC^AgRP-EGFP^ (EGFP, gray) mice during AL or FD conditions. ARC^AgRP-TeLC^ (TeLC, n=9, green) and ARC^AgRP-EGFP^ (EGFP, n=12, gray) mice. (H) Percentage of mice entering torpor bouts (Tb < 33 °C) with a duration longer than an hour (Fisher’s Exact test, odds ratio = 9.14, p = 0.024) (I) Minimum body temperature recorded within each session for ARC^AgRP-TeLC^ (TeLC, green) or ARC^AgRP-EGFP^ (EGFP, gray) mice. Each thin line represents a single animal. Thicker lines and point ranges represent the group. TeLC injections significantly attenuated the fasting-induced temperature drop relative to EGFP (β = 3.14 ± 0.85°C, *t*(19) = 3.71, *p* = 0.001, Satterthwaite d.f.). In the *Ad libitum* fed condition, TeLC and EGFP groups were indistinguishable (0.09 ± 0.70 °C, *t*(35.5) = 0.12, *p* = 0.902). During fasting, EGFP-injected mice showed a drop in minimum body temperature (−5.64 ± 0.55 °C, *p* < 0.001), whereas TeLC-injected mice showed an attenuated response (−2.50 ± 0.64 °C, *t*_₍_19_₎_ = −3.91, *p* < 0.001) (J) Mean torpor bout duration (body temperature < 33 °C for at least 1h) during the fasting session for ARC^AgRP-TeLC^ (TeLC, green) and ARC^AgRP-EGFP^ (EGFP, gray) mice. Each dot is a single animal. TeLC injections reduced mean torpor bout duration (EGFP: 216.92 ± 45.52; TeLC: 49.17 ± 27.74, Wilcoxon’s rank sum test, p = 0.006) Data are represented as Mean ± SEM.

Next, we tested whether ARC^AgRP^ neuronal activity is required for the generation of fasting-induced torpor. For this set of experiments, *Agrp*-IRES-Cre female mice (which are smaller and show longer torpor bouts than males)^16^ were bilaterally injected either with cre-dependent EGFP (ARC^AgRP-EGFP^, control) or TeLC-EYFP (ARC^AgRP-TeLC^), a derivative of tetanus toxin that inhibits synaptic transmission (Fig. 1F). We collected baseline Tb recordings in fed AL mice and tested the ability of these animals to enter torpor during a 24h period of FD starting at lights-off (ZT12) (Fig. 1G). Compared to female controls, ARC^AgRP-TeLC^ mice were less likely to enter long-duration bouts of torpor (Fig. 1H). While minimal Tb in AL fed ARC^AgRP-TeLC^ animals was indistinguishable from AL fed ARC^AgRP-EGFP^ controls, the decrease in Tb in fasted ARC^AgRP-TeLC^ animals showed either a decrease in the depth or the absence of torpor bouts (Fig. 1I and Ext data Fig 2A). In addition, the mean torpor bout duration was decreased from approximately 217 ± 45 minutes in ARC^AgRP-EYFP^ mice to about 49 ± 27 minutes in ARC^AgRP-TeLC^ animals (Fig. 1J). Similarly, the total time spent in torpor was reduced in ARC^AgRP-TeLC^ mice (Ext data Fig 2B,C). Of note, ARC^AgRP-TeLC^ mice showed higher weights compared to controls. However, the observed inhibition of torpor responses was independent of any BW differences (Ext data Fig 2D, E). To exclude the possibility that the effects on torpor were due to off-target Cre-independent expression of TeLC, we injected C57BL/6 mice with AAV-DIO-TeLC. As expected, this experiment showed no changes in the torpor phenotype, suggesting that the attenuation and inhibition of torpor were driven by ARC^Agrp-^specific silencing.

**Fig. 2:**
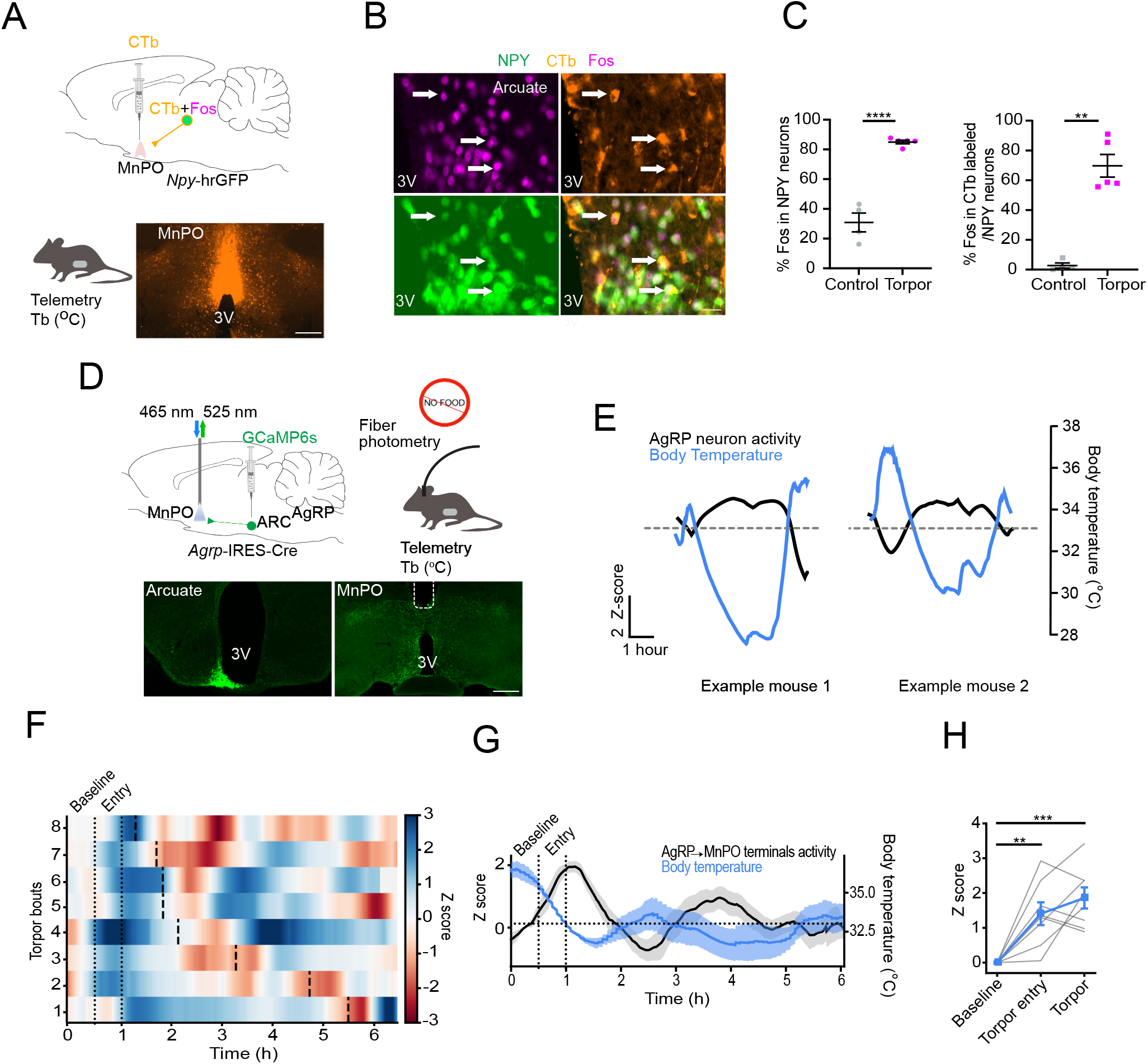
ARC^AgRP^ neurons that innervate the MnPO are activated during torpor. (A) Schematic of approach to map neural afferents to the MnPO that are active during torpor by CTb injection into the MnPO combined with Fos immunostaining (top), in mice implanted with biotelemetry probes to monitor body temperature (Tb) (lower left). Location of CTb injection in the MnPO (lower right). Scale bar = 200 μm (B) Representative images of co-expression of CTb and Fos in AgRP/NPY neurons in the arcuate nucleus. Scale bar = 20 μm (C) Quantification of Fos expression in ARC^NPY+^ neurons of fed AL (control) and food deprived mice (30.9 % ± 6.3 SEM Fos in NPY+ neurons in controls vs 85% ± 1.2 SEM Fos in NPY+ neurons in torpid mice, unpaired t-test p<0.0001, n=4 controls and n=5 torpid mice) and the Fos labelling in CTb/NPY neurons in the arcuate nucleus (2.7% ± 1.7 SEM Fos in CTb/NPY+ neurons in control mice vs 69.7% ± 7.6 SEM Fos in CTb/NPY+ neurons in torpid mice, unpaired t-test p=0.0001, n=4 controls and n=5 torpid mice). (D) Schematic of *in vivo* fiber photometry recordings from ARC^AgRP^→MnPO terminals (top left) concurrent with biotelemetry recordings of Tb (top right) and representative image of GCaMP in ARC^AgRP^ neurons and placement of the optic fiber in the MnPO (lower). Scale bar = 200 μm (E) Representative traces of ARC^AgRP^→MnPO neuron terminals activity and Tb during topor bouts. Dotted horizontal lines: torpor, defined as < 33°C. (F) Heatmap of ARC^AgRP^→MnPO terminal activity during all recorded torpor bouts (n=7 bouts from 3 mice), arranged by bout length. Thick vertical black lines = end of the bout. Grey shaded periods denote bouts where the recording had ended. (G) Mean activity of ARC^AgRP^→MnPO terminals and Tb during torpor bouts. Dotted horizontal lines: topor, defined as < 33°C. (H) Mean activity of ARC^AgRP^→MnPO terminals activity prior to the entry to torpor, during the entry and during torpor bouts. Gray lines: individual mice. One-way repeated measures ANOVA with Tukey’s multiple comparisons test, \*\**p*=0.0021, \*\*\**p*=0.00016. Data are represented as Mean ± SEM.

### The ARC^AgRP→^MnPO pathway is active during fasting-induced torpor

ARC^AgRP^ neurons innervate the MnPO^26^, a canonical center for thermoregulatory control and a region containing a unique group of thermoregulatory neurons required for torpor^17^. We speculated that the ARC^AgRP^→MnPO pathway is active and produces torpor bouts when mice are food deprived. First, as there are no reliable antibodies for AgRP that label cell bodies, and virtually all AgRP neurons express NPY^27^, we used a traditional retrograde tracing method (Cholera toxin subunit B, CTb) in the MnPO of NPY-hrGFP mice and induced fasting-induced torpor in mice that were kept at 22 °C Ta (Fig. 2A). After confirming that mice entered torpor (Tb <33 °C for > 1 hr), we perfused them. Controls were perfused at the same time of the day but were kept with full access to food. Then, we used histological analyses to evaluate whether NPY-expressing ARC (ARC^NPY^) neurons that innervate the MnPO express Fos, a marker for neuronal activation, during torpor. These tests showed that about 85% of all ARC^NPY^ neurons expressed Fos during torpor, and approximately 70% of the ARC^NPY^ cells that project to the MnPO region (CTb+ neurons) were active during fasting-induced torpor (Fig. 2B, C). Hence, the ARC^AgRP^→MnPO pathway is activated during fasting-induced torpor.

Given that not all AgRP neurons project to this preoptic region^28^, we specifically recorded the changes in calcium signaling of AgRP neuron terminals in the MnPO during 24-48 hours of FD condition (Fig. 2D). We found an increase in the GCaMP signal as mice entered torpor (Fig, 2E), and this effect was enhanced during torpor; we observed the highest GCaMP fluorescence levels during the prolonged hypothermic period until Tb reaches its nadir (Fig. 2F). The results of the individual torpor bouts are shown in F and the time-course relationship of Tb and Z-score in G and H. Of note, we also observed decreased GCaMP fluorescence levels at torpor inter-bouts (“arousal” periods) (Ext data Fig 3). The continuous increase in GCaMP signal from ARC^AgRP^ synaptic terminals in the MnPO during torpor and its reduction when mice exit torpor states suggest that this circuit is recruited for torpor regulation.

**Fig. 3.**
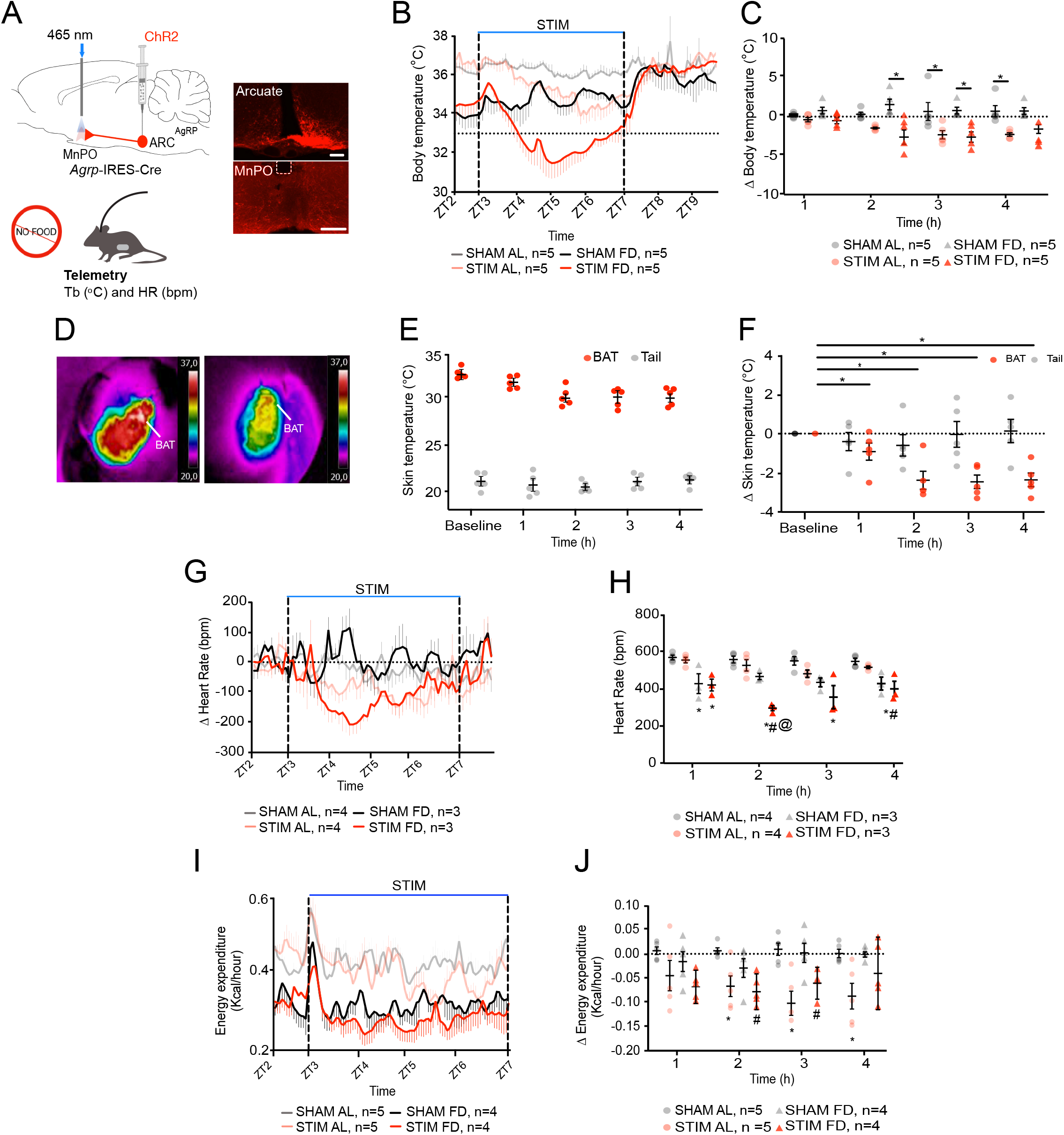
Activation of synaptic terminals from ARC^AgRP^ neurons in the MnPO produces a reduction of core body temperature, heart rate, and energy expenditure. (A) Schematic ChR2-mediated activation of the ARC^AgRP^→MnPO circuit (top left) while monitoring core body temperature (Tb) and heart rate (HR) (lower left). Representative images of ChR2 expression in AgRP-expressing neurons in the arcuate nucleus and their synaptic terminals in the MnPO below the area of the fiber placement (right). Scale bar 100 μm (B) Mean body temperature recorded from *AgRP*-cre mice from ZT2-ZT9, ChR2-mediated activation of the ARC^AgRP^ →MnPO pathway for 4-h (ZT3-ZT7) caused hypothermic responses in animals fed *ad libitum* (AL, n=5) (Mean Tb ZT3-ZT7: 35.8 ± 0.09 °C SHAM AL vs. 33.9 ± 0.12 °C STIM AL, paired t-test, p<0.0001) or food deprived (FD, n=5) mice (Mean Tb ZT3-ZT7: 34.2 ± 0.1 °C SHAM FD vs. 32.8 ± 0.15 °C STIM FD, paired t-test, p<0.0001). (C) Quantification of the hypothermic responses of mice subjected to activation of the ARC^AgRP^→MnPO (STIM) or controls (SHAM) during AL or FD conditions (Mean Tb change in an hourly basis: 1-h: 0.17 ± 0.14 °C SHAM AL vs. -0.39 ± 0.29 °C STIM AL, 0.77 ± 0.42 °C SHAM FD vs. -0.5 ± 0.39 °C STIM FD; 2-h: 0.3 ± 0.27 °C SHAM AL vs. -1.46 ± 0.13 °C STIM AL, 1.55 ± 0.68 °C SHAM FD vs. -2.62 ± 0.92 °C STIM FD; 3-h: 0.65 ± 1.16 °C SHAM AL vs. -2.32 ± 0.54 °C STIM AL, 0.77 ± 0.42 °C SHAM FD vs. -2.62 ± 0.66 °C STIM FD; 4-h: 0.70 ± 0.73 °C SHAM AL vs. -2.28 ± 0.23 °C STIM AL, 0.71 ± 0.43 °C SHAM FD vs. -1.59 ± 0.56 °C STIM FD two-way ANOVA, followed by Bonferroni’s post hoc test, *^#^p<0.05, n=5 each group). (D) Representative infrared images from before and after ARC^AgRP^→MnPO stimulation. (E) Temperature of brown adipose tissue (BAT) and tail skin before and after ARC^AgRP^→MnPO stimulation. (F) Quantification of the changes in skin temperature above the brown adipose tissue (BAT) and tail temperatures shows a reduction in BAT temperature 2, 3, and 4 h from the stimulation onset (Difference in BAT temperature relative to baseline: -0.92 ± 0.43 1H STIM AL vs. -2.3 ± 0.46 2H* STIM AL vs. -2.46 ± 0.034 3H* STIM AL vs. - 2.36 ± 0.34 4H* STIM AL, two-way ANOVA, followed by Bonferroni’s post hoc test, * p<0.0001, n=5). (G) Mean change in heart rate (HR) in response to SHAM or activation of the ARC^AgRP^→ MnPO pathway during AL (n=4) or FD (n=3) conditions. (H) Quantification of the HR in mice during stimulation of the ARC^AgRP^→MnPO pathway (STIM) or controls (SHAM) during AL and FD conditions. Mean HR 1h: 569.3 ± 12.65 bpm SHAM AL vs. 428.4 ± 53.27 bpm SHAM FD or 421.0 ± 31.69 bpm STIM FD, One-way ANOVA followed by Bonferroni’s post hoc test, * p < 0.05, and 555.3 ± 15.11 bpm STIM AL vs. 421.0 ± 31.69 bpm STIM FD, One-way ANOVA followed by Bonferroni’s post hoc test, * p=0.04. Mean HR 2h: 558.2 ± 18.55 bpm SHAM AL vs. 296.5 ± 13.87 bpm STIM FD, One-way ANOVA followed by Bonferroni’s post hoc test, **** p<0.0001, 527.2 ± 30.52 bpm STIM AL vs. 296.5 ± 13.87 bpm STIM FD, One-way ANOVA followed by Bonferroni’s post hoc test, *** p=0.0002, and 466.9 ± 16.93 bpm SHAM FD vs 296.5 ± 13.87 bpm STIM FD, ** p=0.003. Mean HR 3h: 551.0 ± 22.79 bpm SHAM AL vs. 355.7 ± 62.24 bpm STIM FD, One-way ANOVA followed by Bonferroni’s post hoc test, * p=0.01. Mean HR 4h: 548.4 ± 16.67 bpm SHAM AL vs. 400.5 ± 40.9 bpm STIM FD, One-way ANOVA followed by Bonferroni’s post hoc test, ** p=0.01, and 516.5 ± 7.0 bpm STIM AL vs. 400.5 ± 40.9 bpm STIM FD, One-way ANOVA followed by Bonferroni’s post hoc test, * p=0.04. (I) Mean energy expenditure from ZT2-ZT9, ChR2-mediated activation of the ARC^AgRP^→MnPO pathway for 4-h (ZT3-ZT7) in mice fed *ad libitum*. (J) Quantification of the change in energy expenditure of mice subjected to activation of the ARC^AgRP^→MnPO (STIM) or controls (SHAM) during AL or FD conditions. Mean change EE for mice kept AL, 2H: 0.006 ± 0.006 kcal/hr SHAM AL vs. -0.06 ± 0.022 Kcal/hr STIM AL, paired t-test, * p=0.019; 3H: 0.01 ± 0.01 kcal/hr SHAM AL vs. - 0.102 ± 0.025 Kcal/hr STIM AL, paired t-test, * p=0.02; 4H: 0.001 ± 0.009 kcal/hr SHAM AL vs. -0.08 ± 0.02 Kcal/hr STIM AL, paired t-test, ** p=0.008, n=5. In mice subjected to FD, the mean change in EE 2H: -0.029 ± 0.019 kcal/hr SHAM FD vs. - 0.078 ± 0.013 Kcal/hr STIM FD, paired t-test, *p=0.03 and 3H: 0.002 ± 0.02 kcal/hr SHAM FD vs. -0.061 ± 0.012 Kcal/hr STIM FD, paired t-test, **p=0.007, n=5. Data are represented as Mean ± SEM.

### Activating the ARC^AgRP^→MnPO pathway induces a decrease in body temperature, heart rate and energy expenditure

Torpor is characterized by a regulated hypothermic response in which animals switch off heat production against cold signals, reducing their thermoregulatory energy costs and EE. This survival mechanism enables energy conservation in conditions of food scarcity in energetically demanding environments (e.g., cold). To further test the role of the ARC^AgRP^→MnPO neural pathway in producing the adaptive physiological responses typical of torpor, we injected Cre-dependent Channelrhodopsin (ChR2) into the ARC of *Agrp*-IRES-cre male mice, while optical fibers were implanted over the MnPO (Fig. 3A). Then, we optogenetically activated the ARC^AgRP^→MnPO circuit for 4 h during the light period (ZT3-ZT7) while monitoring the changes in Tb, HR, EE, and the indirect temperature of the brown adipose tissue (BAT).

We tested the effects of activating the ARC^AgRP^→MnPO pathway in freely moving mice kept at 22 °C Ta in both fed AL or FD conditions. Our experiments demonstrated that activating the ARC^AgRP^→MnPO pathway caused a decrease in Tb of up to 2.8 °C in fed AL mice and 3.1 °C in the same fasted animals (paired t-test, p<0.0001, n=5) (Fig. 3B, C). Moreover, using infrared imaging, we discovered that BAT thermogenesis is suppressed upon stimulation of the ARC^AgRP^→MnPO pathway (Fig. 3D-F). On the other hand, activation of the ARC^AgRP^→MnPO circuit did not produce changes in the tail temperature of mice, suggesting that heat dissipation is not involved in the reduction of Tb caused by the stimulation of this pathway. These results indicate that suppressed BAT thermogenesis, but not tail vasodilation, drives the reduction in Tb that we observed when this pathway is activated.

To further investigate the effects of activating the ARC^AgRP^→MnPO circuit, we measured changes in HR, finding a reduced HR during activation of the ARC^AgRP^→MnPO circuit in AL or FD conditions (Fig. 3G,H). Next, we measured EE using indirect calorimetry. Activating the ARC^AgRP^ synaptic terminals in the MnPO reduced EE of mice when they were given full access to food or not (Fig. 3I,J), which suggests that activation of the ARC^AgRP^→MnPO pathway mimics physiological responses typically observed in mice during fasting-induced torpor bouts. The decrease of EE was slightly reduced during the stimulation of the ARC^AgRP^→MnPO pathway when animals were food restricted. We speculate that the AgRP neurons are already activated in fasted mice, and stimulating them further may have a modest effect.

While activation of the ARC^AgRP^→MnPO circuit was sufficient to produce hypothermia and reduce EE, photostimulating this pathway in mice fed AL did not induce a sustained reduction of Tb, mimicking the degree and length typically observed during spontaneous fasting-induced torpor bouts in laboratory mice^1,24,29^. Given that activating ARC^AgRP^ neurons causes mice to eat, we assessed whether stimulating the ARC^AgRP^ neuron projection induces feeding. We found that activation of this circuit increased food intake in a subset of mice (Ext data Fig 4A-C), suggesting that the delayed and reduced degree of hypothermia in mice with unlimited access to food may at least in part be caused by the Tb prandial effect due to food consumption.

### Inhibiting the ARC^AgRP^→MnPO pathway prevents fasting-induced torpor

We also used an optogenetic method to inhibit the ARC^AgRP^→MnPO neural pathway (Fig. 4A) to test whether this pathway was necessary for torpor induction. We injected Cre-dependent *eOPN3*-*mScarlet*-(eOPN3) into the ARC of *Agrp*-IRES-Cre mice and used low power illumination through optical fibers implanted over the MnPO to cause a long-lasting inhibition of synaptic release from ARC^AgRP^ synaptic terminals in the MnPO. Then, we inhibited this pathway for 18 h starting at the lights-off period (ZT12-ZT6), when we also initiated our food deprivation protocol. We found that inhibiting AgRP synaptic terminals in the MnPO entirely prevented fasting-induced torpor in these mice (Fig. 4B, C), without altering locomotor activity patterns during the period when they normally display torpor bouts (Ext Fig 5A-B). Additionally, we used animals with a missed injection of eOPN3 or a misplaced optical fiber as controls. These mice displayed similar torpor responses during FD associated with SHAM or light stimulation (Ext Fig 5C,D).

**Fig. 4:**
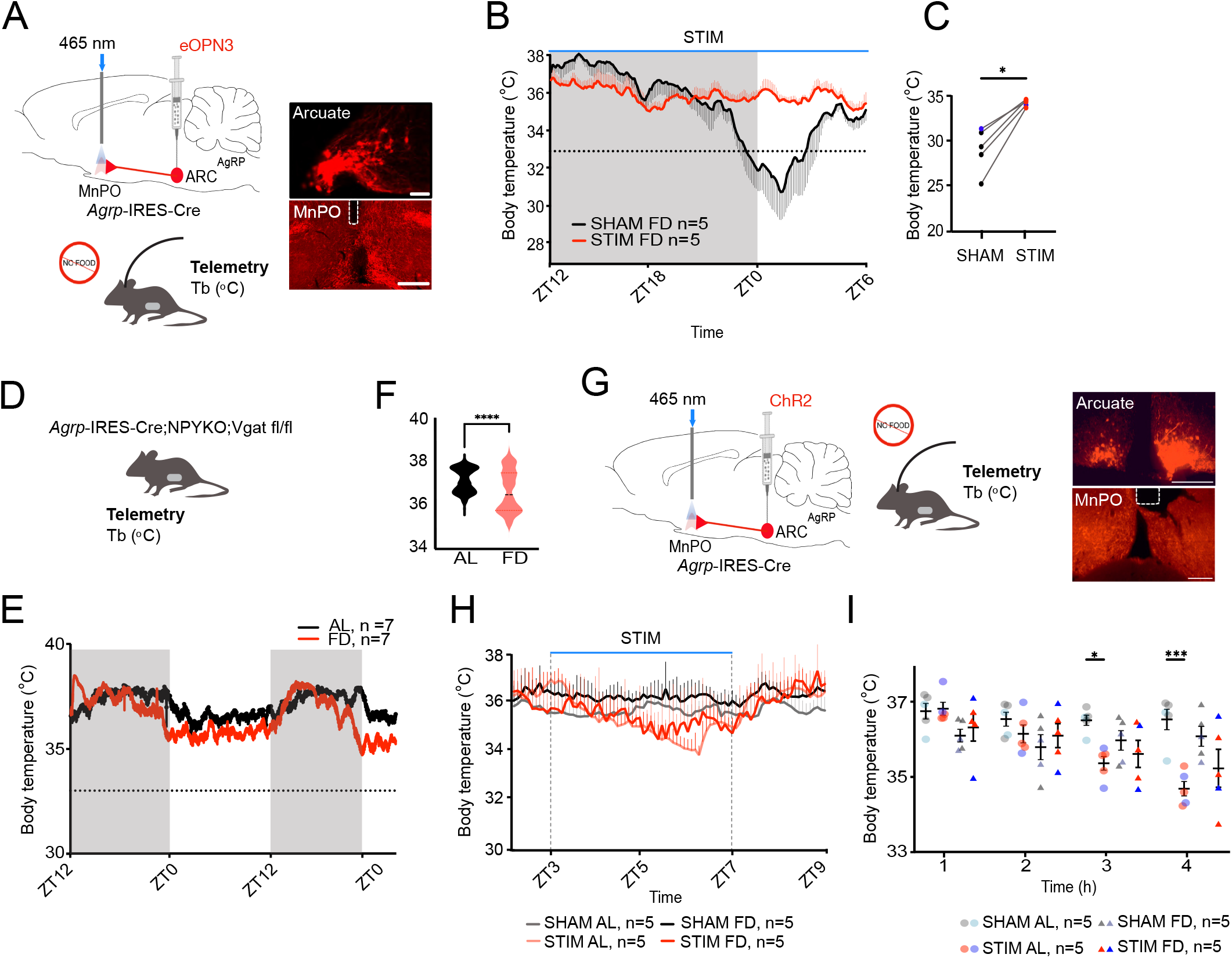
The ARC^AgRP^→MnPO pathway is required for fasting-induced torpor. (A) Schematic of optogenetic inhibition of the ARC^AgRP^→MnPO circuit (upper left) while monitoring core body temperature (Tb) of mice (lower). Representative images of AAV-eOPN3 expression (red) in AgRP-expressing neurons in the arcuate nucleus (upperright) and their synaptic terminals in the MnPO below the area of the fiber placement (bottom right). (B) Mean body temperature of food deprived (FD, n= 4 females and 1 male) *AgRP*-cre mice, which was recorded for 18-h from ZT12 to the next day (ZT6) during SHAM or photoinhibition of the ARC^AgRP^→MnPO pathway. (C) Quantification of the mean minimal body temperature 29.24 ± 1.1 °C SHAM FD vs. 34.4 ± 0.17 °C STIM FD, paired t-test, **p=0.005 (n=5, 4 females are reported in red and 1 male mouse in blue). (D) A schematic of the recording of body temperature from *AgRP*-cre;NPYKO;Vgatfl/fl mice using biotelemetry. (E) Changes in body temperature of *AgRP*-cre;NPYKO;Vgatfl/fl mice subjected to FD or fed *ad libitum* (baseline) conditions during 48 hours. (F) Quantification of mean Tb of *AgRP*-cre;NPYKO;Vgatfl/fl mice during a 48 h period: 37.13 ± 0.02 °C Baseline vs. 36.64 ± 0.04 °C FD, paired t-test, p<0.0001, n= 4 females and 3 males. (G) Schematic of optogenetic activation of the ARC^AgRP^→MnPO circuit (left)while monitoring core body temperature (Tb) (center) of *AgRP*-cre;NPYKO;Vgatfl/fl mice (top). Representative images of AAV-ChR2 expression (red) in AgRP-expressing neurons in the arcuate nucleus (upper right) and their synaptic terminals in the MnPO (lower right), which is below the area of the fiber placement, in a *AgRP*-cre;NPYKO;Vgatfl/fl mouse. (H) Mean body temperature recorded from *AgRP*-cre;NPYKO;Vgatfl/fl mice during activation of the ARC^AgRP^→MnPO (STIM) or controls (SHAM) in mice fed ad libitum (AL), or the same mice subjected to FD. (I) ChR2-mediated activation of the ARC^AgRP^ →MnPO pathway for 4-h (ZT3-ZT7) caused a small reduction of Tb in AL condition (Mean Tb 3h: 36.64 ± 0.07 °C SHAM AL vs. 35.91 ± 1.0 °C STIM AL, two-way ANOVA, followed by Bonferroni’s post hoc test, *p<0.05, n=5; Mean Tb 4h: 36.63 ± 0.07 °C SHAM AL vs. 36.16 ± 0.4 °C STIM AL, two-way ANOVA, followed by Bonferroni’s post hoc test, * p<0.0001, n=5). The individual data is shown, n = 3 females (grey SHAM and red STIM) and 2 males (light blue SHAM and dark blue STIM). Data are represented as Mean ± SEM.

Then, we dissected the synaptic mechanism by which ARC^AgRP^ neurons produce adaptive responses that suppress energy expenditure and lower Tb for bridging periods when energy supplies are limited. AgRP neurons release the inhibitory neurotransmitters GABA, NPY and AgRP. We first used transgenic mice that are unable to release GABA from AgRP neurons due to selective removal of the vesicular GABA transporter, *Slc32a1* (vGAT) (*Agrp*-IRES-cre;Vgat^fl/fl^ mice). We fasted these mice for 24 h (Ext Fig 5E), which started at the lights off (ZT12), and observed that all *Agrp*-IRES-cre;Vgat^fl/fl^ mice tested under FD showed torpor responses (Ext data Fig 5F). Then, we optogenetically activated the ARC^AgRP^→MnPO circuit in *Agrp*-IRES-cre;Vgat^fl/fl^ mice (Ext data Fig 6A). For this, we injected Cre-dependent ChR2 into the ARC of these mice, while optical fibers were implanted over the MnPO. Using a 4-h light stimulation period, we observed a strong Tb reduction in response to the activation of the ARC^AgRP^→MnPO pathway (Ext data Fig 6B-D). Hence, GABA alone does not appear necessary for fasting-induced torpor or for the hypometabolic and hypothermic state caused by ARC^AgRP^→MnPO activation.

Then, we tested whether the release of NPY from ARC^AgRP^ neurons enables hypothermic responses during total FD by using mice that globally lack NPY (*Agrp*-IRES-cre;NPYKO mice). Similar to a previous report^8^, we found that the impaired release of NPY was insufficient to fully preclude the drop of Tb or torpor responses in all tested animals. However, two out of four *Agrp*-IRES-cre;NPYKO mice did not show Tb below 33 °C for at least one hour during the period of FD (Ext data Fig 5F), which is typical for fasting-induced torpor bouts^17^. As some *AgRP*-cre;NPYKO mice failed to produce fasting-induced torpor responses, we wanted to further dissect the role of NPY in producing hypothermic responses. For this, we first infused NPY (4 μg in 1 μl) directly into the MnPO of mice, which were restricted of food access for 5 h - starting 2 h before the single NPY injection. We found a robust NPY-induced hypothermic effect that lasted for about 3 hours - the period in which the animals did not have access to food (Ext data Fig 7A, B). The hypothermic effect of NPY in the POA suggests a critical role for this neuromodulator in Tb control, and because this response has been previously reported in rats^30^, it implies that NPY-mediated action on preoptic neurons is conserved across these mammalian species. On the other hand, when using ChR2-mediated activation of the ARC^AgRP→^MnPO pathway in *AgRP*-cre;NPYKO mice (Ext data Fig 7C-F), these animals produced hypothermic responses. Together, these results indicate that other neurotransmitters likely act in synergy with NPY to produce hypothermic responses, including fasting-induced torpor.

Next, we used mice lacking the ability to release both GABA and NPY (*Agrp*-IRES-cre; NPYKO; Vgat^fl/fl^ mice), and found that these animals never entered torpor during a 48 h period of FD. Evaluating the changes in Tb of these triple transgenic mice (*Agrp*-IRES-cre; NPYKO; Vgat^fl/fl^ mice), we observed a slight decrease in their Tb (∼ 0.5 °C) during the 48-h FD protocol (Fig. 4D-F). We also tested the contribution of NPY and GABA to the hypothermic response produced by stimulation of the ARC^AgRP^→MnPO pathway. For this, we repeated the ChR2-mediated circuit stimulation of the ARC^AgRP→^MnPO pathway (see Fig 3B, which caused hypothermic responses with an Tb decrease of about 3 °C in both AL and FD conditions in intact mice) in *Agrp*-IRES-cre; NPYKO;Vgat^fl/fl^ mice for 4 h (ZT3-ZT7). This optogenetic approach modestly reduced Tb (∼0.6 °C) in these double KO mice under both the AL and FD conditions (Fig. 4G-I). To test whether local heat production in the POA due to laser stimulation triggered the Tb reduction effect observed in the *Agrp*-IRES-cre; NPYKO;Vgat^fl/fl^ mice, we injected additional mice with AAV-mCherry. These controls did not show Tb reduction in response to our optogenetic stimulation (Ext data Fig 7G-J). This suggests that neuromodulators other than NPY and GABA (e.g., AgRP) may drive the small reduction of Tb observed when activating the ARC^AgRP^→MnPO pathway in *Agrp*-IRES-cre; NPYKO;Vgat^fl/fl^ mice. However, these mediators appear to play only a modest thermoregulatory role. Indeed, our results indicate that they are not necessary for torpor response. Thus, our data support the idea that the co-release of NPY and GABA inhibitory neurotransmitters from ARC^AgRP^ neurons is required for fasting-induced torpor.

### MnPO^EP3R^ neurons mediate the decrease in body temperature caused by activation of the ARC^AgRP^→MnPO pathway

We next tested whether activating the ARC^AgRP^→MnPO pathway causes an increase in the activity of MnPO^EP3R^ neurons. For this, we activated the synaptic terminals of ARC^AgRP^ neurons in the MnPO using the red-shifted opsin ChrimsonR, while simultaneously recording Tb and changes in calcium signaling in MnPO^EP3R^ neurons using fiber photometry (Fig. 5A). Optogenetic stimulation of the ARC^AgRP^→MnPO neural pathway for 2 h during the light (starting at ZT13) or dark (starting at ZT15) phases caused an increase in intracellular calcium levels in the MnPO^EP3R^ neurons, while causing a decrease in Tb (Fig. 5B-D) of a similar magnitude to our ChR2-mediated stimulation of the ARC^AgRP^→MnPO pathway (Fig. 3). Next, we developed a new AAV to express Cre recombinase under the promoter for *Ptger3* (the gene encoding EP3R) (Ext data Fig 8A), and co-inject this AAV-cre_Ptger3 with the cre-dependent AAV-hM3Dq to allow chemogenetic activation of the EP3R-expressing MnPO neurons (Ext data Fig 8B, C). With this method, we observed a reduction in core Tb from ∼37 °C to ∼28 °C, replicating the hypothermic profile identified in our previous studies^17^ and during chemogenetic activation of MnPO^EP3R^ neurons in transgenic Ptger3^Cre^ mice (Ext data Fig 8D). We also used this new AAV to ablate the MnPO^EP3R^ neurons. For this, we used an AAV mix of AAV-cre_Ptger3 and AAV-DIO-caspase in the MnPO of AgRP-cre mice. These same animals were also bilaterally injected with Cre-dependent ChR2 into the ARC to enable optogenetic activation of the ARC^AgRP^→MnPO pathway. We found that the hypothermic response caused by activation of ARC^AgRP^ synaptic terminals in the MnPO was attenuated in mice lacking MnPO^EP3R^ neurons (a maximum Tb decrease of ∼ 2.3 °C in intact animals compared to ∼1.5 °C in ablated mice), suggesting that the hypothermic response observed during torpor bouts may be mediated by the ARC^AgRP^→MnPO^EP3R^ pathway (Ext data Fig 8E-H).

**Fig. 5:**
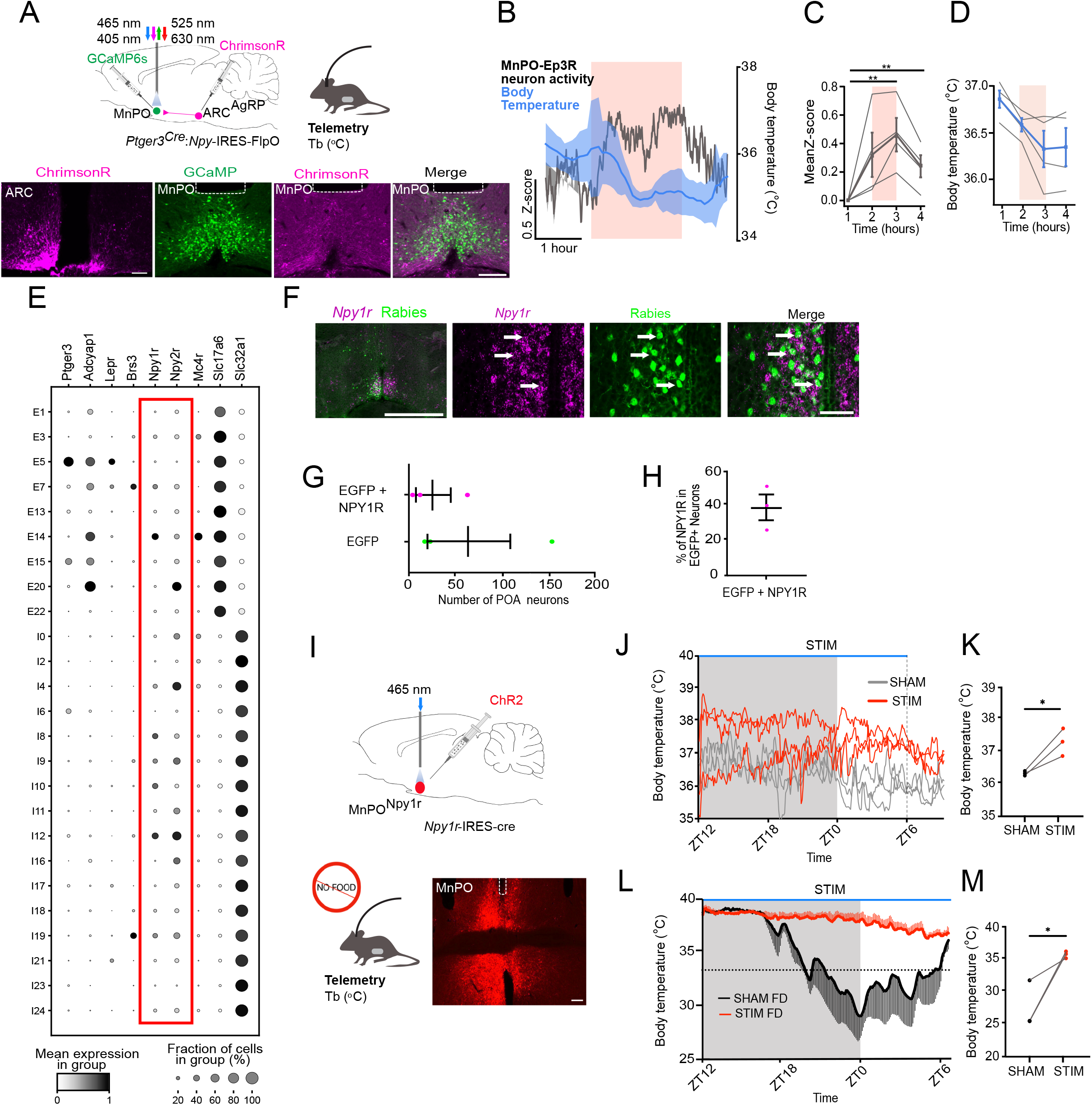
Activation of the ARC^AgRP^→MnPO pathway produces an increase in calcium signaling in MnPO^EP3R^ cells, which is associated with a reduction of core body temperature. (A) Schematic of *in vivo* fiber photometry recordings from MnPO^EP3R^ neurons with concurrent with optogenetic stimulation of ARC^AgRP^ neuron terminals in the MnPO using ChrimsonR (upper left), along with biotelemetry recordings of Tb (upper right) and a representative image of ChrimsonR in ARC^AgRP^ neurons (lower left) and placement of the optic fiber in the arcuate (lower right). Scale bar = 200 μm. (B) Mean activity of MnPO^EP3R^ neurons and core body temperature during optogenetic stimulation of ARC^AgRP^ neuron terminals in the MnPO (5-6 trials per animal during the light and dark phases). The period of optogenetic stimulation is denoted by the pink bar (n = 4). (C) Mean activity of MnPO^EP3R^ neurons prior to, during and after stimulation of ARC^AgRP^ neuron terminals in the MnPO. Gray lines: individual mice (n = 4). One-way ANOVA with Tukey’s multiple comparisons test, ** *p*=0.004 (D) Mean body temperature of mice prior to, during, and after stimulation of ARC^AgRP^ neuron terminals in the MnPO: 1h: 36.859 ± 0.093 °C, 2h: 36.585 ± 0.072 °C, 3h: 36.325 ± 0.195 °C, 4h: 36.346 ± 0.205 °C. Gray lines: individual mice (n = 4). Paired t-test, * p=0.03. (E) Dot plot depicting the mean gene expression (minmax-normalized by gene, grayscale) and the fraction of cells identified in each cluster expressing each gene (size of circles). Neuronal cell type labels are based on clustering in Ext data Fig. 9, and comprise neuronal class (E, excitatory; I, inhibitory), followed by cluster number. (F) Representative images of monosynaptic rabies tracing in the POA with colocalization of *Npy1r* mRNA. (G) Number of POA neurons that express rabies-EGFP and *Npy1r* mRNA (magenta) or rabies alone (green). Each dot is an individual animal (n =3). (H) Quantification of % of rabies-EGFP neurons that express *Npy1r* mRNA. Each dot is an individual animal (n=3). (I) Schematic of approach using ChR2-mediated optogenetic activation of the POA^Npy1r^ (top) while monitoring core body temperature (Tb) (lower left) of *Npy1r-*IRES-Cre mice (top). Representative images of AAV-ChR2 expression (red) in Npy1r-expressing neurons and fiber placement in the MnPO (lower right).Mean body temperature recorded from *Npy1r*-cre mice during AL or FD conditions. (J) ChR2-mediated activation of the POA^Npy1r^ neurons (ZT12-ZT6) caused an increase in body temperature. (K) Mean change in body temperature before and after ChR2-mediated activation of the POA^Npy1r^ neurons in mice fed AL: 36.33 ± 0.04 °C SHAM vs. 37.3 ± 0.22 °C STIM, paired t-test, p = 0.04, n = 2 males and 1 female. (L) ChR2-mediated activation of the POA^Npy1r^ neurons prevented torpor. (M) Minimal body temperature before and after ChR2-mediated activation of the POA^Npy1r^ neurons during the FD condition: 27.55 ± 2.0 °C SHAM FD vs. 35.62 ± 0.3 °C STIM FD, paired t-test, p = 0.03, n = 2 males and 1 female. For the individual traces see Ext data Fig 11A,B. Data are represented as Mean ± SEM.

ARC^AgRP^ neurons are inhibitory as they release only inhibitory neurotransmitters. Hence, for ARC^AgRP^ neurons to activate MnPO^EP3R^ neurons, they must do so by disinhibition. This could be achieved by inhibiting GABAergic neurons that inhibit MnPO^EP3R^ neurons under conditions in which mice have sufficient energy stores and/or access to food sources, or by inhibiting the axon terminals of GABAergic afferents that provide inhibitory tone to MnPO^EP3R^ neurons. The increased activity of ARC^AgRP^ neurons would remove this inhibitory tone, releasing the activity of MnPO^EP3R^ neurons.

### Activating NPY1R and NPY2R-expressing POA neurons prevents torpor

As our studies identified NPY as a key neuromodulator used by ARC^AgRP^ cells to induce torpor, we hypothesized that preoptic neurons expressing NPY receptors are likely the direct targets of ARC^AgRP^ cells, functioning as inhibitory interneurons in the ARC^AgRP^→MnPO^EP3R^ neural pathway. To identify which neuronal populations express NPY receptors in the POA, we reanalyzed a spatial transcriptomic dataset from the Allen Brain Institute^31^. Among the 16 GABAergic and 9 glutamatergic neuronal classes in the POA (Fig 5E and Ext data Fig 9A-G), we found that the *Npy1r* and *Npy2r* genes are expressed in several neuronal populations, but not in the neurons that express *Ptger3*.

Studies using Siberian hamsters have identified the NPY1 receptor (NPY1R) as a brain mediator of NPY-induced torpor-like hypothermia^32–34^. However, to our knowledge, the specific location of the NPY1R-expressing neurons required to generate hypothermic responses has not been revealed by previous work. We speculated that NPY1R-expressing POA (POA^Npy1r^) neurons would be required for the generation of torpor. In our model, POA^Npy1r^ cells may tonically inhibit MnPO^EP3R^ thermoregulatory neurons during periods of normal energy balance, and the release of NPY and GABA from ARC^AgRP^ neurons inhibits POA^Npy1r^ neurons, thereby disinhibiting MnPO^EP3R^ neurons and enabling animals to enter bouts of torpor only when energy stores are low. In line with this, using monosynaptic tracing combined with *in situ hybridization* (ISH), we found robust connectivity between Npy1r-expressing preoptic neurons and the MnPO^EP3R^ neurons (Fig 5F-H), and no evidence that ARC^AgRP^ neurons directly innervate MnPO^EP3R^ cells (Ext data Fig 10A-C). As many POA^Npy1r^ neurons are GABAergic and they directly innervate the MnPO^EP3R^ neurons, we speculated that activating the POA^Npy1r^ neurons would prevent the induction of torpor.

To dissect the physiological role of POA^Npy1r^ neurons in the brain circuit for torpor, we optogenetically activated these neurons in AL fed and fasted female and male mice (Fig. 5I and Ext data Fig. 11A, B). For this, we injected Cre-dependent ChR2 and implanted optical fibers into the POA of *Npy1r*^cre^ mice. Using the same method, we also tested the role of the Npy2r-expressing POA neurons in torpor (Ext data Fig 11C). We activated the POA^Npy1r^ and POA^Npy2r^ neurons during an 18-h period (ZT12-ZT6) in mice that had free access to food or food deprived animals (food restriction also started at ZT12). When POA^Npy1r^ neurons were activated in AL fed mice, there was an increase in core Tb of about 1 degree Celsius (Fig. 5J,K). Under the FD condition, activating POA^Npy1r^ neurons fully prevented mice from producing a torpor response (Fig. 5L,M). Thus, the effect of activating POA^Npy1r^ neurons in mice that did not have access to food was significantly greater than the hyperthermic effect observed in satiated animals. Although activation of the POA^Npy2r^ neurons prevented torpor in some animals, it also produced a Tb reduction during the first hours of the optogenetic stimulation (Ext data Fig 11D, E) in most mice. The dual effect of POA^Npy2r^ neurons over Tb is likely due to the more heterogeneous nature of the POA^Npy2r^ neural population compared to the POA^Npy1r^ cells. Together, our results suggest that NPY1R and NPY2R are essential for NPY signaling and Tb effects in the POA (Ext data Fig 12), which causes hypothermia and torpor.

## Discussion

Homeostatic mechanisms regulated by the central nervous system maintain Tb of mammals in a constant range, with daily rhythms underlying small Tb fluctuations due to various physiological and behavioral activities over the course of the day. However, to enable survival under adverse conditions, adaptive responses must produce changes in thermoregulation, bringing Tb to higher (e.g., inflammatory fever) or lower regulated levels. Here, we revealed a brain circuit for body cooling in response to low energy stores. We dissected how information about energy deficit is relayed from ARC^AgRP^ cells to neurons in the MnPO region, a brain center for thermoregulatory control that contains neurons necessary for fasting-induced torpor^14,15,17,19,35,36^. First, we evaluated whether ARC^AgRP^ neurons are required for torpor bouts. We found that ARC^AgRP^ neurons are active during torpor bouts, and silencing these neurons impairs torpor generation. We also found that topor-active ARC^AgRP^ cells project to the MnPO, and optogenetically activating the ARC^AgRP^→MnPO pathway reduces Tb, HR and EE, and suppresses BAT thermogenesis, which are physiological mechanisms that enable daily torpor. Next, we used an optogenetic approach to test the necessity of the ARC^AgRP^→MnPO pathway to generate fasting-induced torpor, and found that inhibiting ARC^AgRP^ synaptic terminals in the MnPO abolished torpor bouts. To further characterize the mechanisms by which this hypothalamic pathway enables torpor, we studied the neuromodulators used by ARC^AgRP^ neurons to produce Tb reduction. We found that the co-release of both NPY and GABA from ARC^AgRP^ neurons is necessary for the full hypothermic response associated with the fasted state or during activation of the ARC^AgRP^→MnPO pathway. Moreover, we show that the circuit underlying torpor depends on NPY1R signaling.

In response to starvation, turning off heat production (e.g., BAT thermogenesis) to cause Tb reduction is an effective energy-conserving mechanism^1^. ARC neurons have been previously implicated in the generation of fasting-induced torpor in mice^8,9^. However, the identity of the ARC cells involved in the complex torpor phenomenon and the brain circuits that trigger the reduction in Tb and energy conservation in response to food restriction remain unknown. ARC^AgRP^ neurons are activated by caloric deficit and drive a variety of behavioral and physiological responses related to the fasted state, including increased food intake, stimulation of corticosterone release, and reduced EE^12,13,23,37,38^. We hypothesized that the increase in ARC^AgRP^ neuronal activity in response to food deprivation is also necessary to trigger the accompanying reduction in Tb in a cool environment.

Using fiber photometry and a cell-specific method to block synaptic transmission from ARC^AgRP^ neurons, we demonstrated that these neurons are not only recruited but also necessary for the generation of torpor bouts. However, we found discrepancies in the calcium signaling dynamics of ARC^AgRP^ cell bodies (“active” during torpor, but not at the torpor entry) when compared to their synaptic terminals in the MnPO, which showed a progressive increase in calcium signaling from torpor entry through the deepest torpor states, coinciding with the nadir in core body temperature. As subgroups of ARC^AgRP^ neurons have previously been identified by Campbell and colleagues^39^, we speculate that they may respond differently during torpor. It is possible that the changes in calcium signaling of the torpor-active ARC^AgRP^ cell bodies were masked by another subset of AgRP+ neurons during torpor entry, which presumably would not be involved with torpor induction and may not project to the MnPO.

Activating the ARC^AgRP^→MnPO circuit is sufficient to induce a hypothermic response and states of energy conservation, but we speculate that the torpor regulation may involve additional brain circuits and mechanisms. For example, the circadian clock influences the timing of torpor bouts^24^, suggesting that centers for circadian control may provide additional inputs to MnPO torpor neurons^40^. Nevertheless, as fasting is a prerequisite for torpor in mice, we propose that the ARC^AgRP^→MnPO pathway is required for torpor generation. Because AgRP neurons release only inhibitory neurotransmitters, we hypothesize that activation of ARC^AgRP^ neurons causes the disinhibition of torpor-driving neurons in the MnPO. Further supporting this, we have (1) recently shown that MnPO^GABA^ neurons have a reduction in their intracellular calcium levels during spontaneous fasting-induced torpor bouts^17^, and (2) we now show that activating the ARC^AgRP^ afferentes in the MnPO causes an increase in calcium signaling in MnPO^EP3R^ neurons, which are glutamatergic neurons that produce torpor-like responses when activated^17^. Moreover, ablation of MnPO^EP3R^ neurons attenuated the hypothermic response induced by activation of the ARC^AgRP^→MnPO pathway. This mechanism would be similar to what has been previously demonstrated, that AgRP neurons activate another adaptation to the fasted state by a disinhibitory synaptic mechanism-activation of the HPA axis. Specifically, ARC^AgRP^ neurons projecting to the PVH activate corticotropin-releasing hormone neurons (PVH*^Crh^* neurons) by inhibiting the terminals of GABAergic projections that otherwise constrain the activity of the PVH*^Crh^* neurons and thus, activation of the HPA axis^13^. Together with our current work, this suggests that ARC^AgRP^ neurons exert several of their powerful effects on behavior and metabolism via an indirect synaptic mechanism.

In addition, we propose that some of the GABAergic cells act to prevent torpor when mice are satiated and have appropriate energy stores express NPY receptors. This proposed model is supported by our analysis of the genes expressed by POA neurons^31^, which revealed that POA^Npy1r^ and POA^Npy2r^ neurons mostly belong to inhibitory clusters, and our monosynaptic tracing experiments showing that NPY1R-expressing POA neurons, but not ARC^AgRP^ neurons, directly innervate MnPO^EP3R^ “torpor” neurons. We also demonstrated that, when activated, POA^Npy1r^ increases Tb and prevents fasting-induced torpor. In some cases, activating POA^Npy2r^ neurons prevented mice from entering torpor. We therefore propose that a population of POA inhibitory interneurons, which acts to prevent torpor when energy stores and food availability are sufficient, expresses NPY1R. We also speculate that co-expression of NPY1R and NPY2R may identify a subset of the POA^Npy1r^ inhibitory interneurons that prevent torpor generation.

Small mammals have a larger body surface area relative to body mass, which enables more efficient heat loss^41,42^. Mice weigh roughly 3,000 times less than adult humans and, far more than large mammals, robustly activate BAT thermogenesis when kept in cold conditions. This size-dependent difference in heat loss and thermogenic capacity is also found in humans: infants have a comparatively larger surface area relative to body mass and retain more BAT than adults, yet still show greater susceptibility to hypothermia. Following this size-dependent logic, adult humans would be expected to develop only mild hypothermia in response to low energy stores, even when exposed to a cold environment, which is consistent with clinical reports of hypothermia in severely calorically restricted individuals^43^. Indeed, we expect that the ARC^AgRP^→MnPO circuit that promotes energy conservation is preserved across species, since caloric restriction induces hypometabolic states in several mammalian species, including humans^4,5,43^. Hence, understanding the homeostatic and adaptive mechanisms that mediate the interaction between energy demands and homeothermy can be leveraged to identify central targets for modulating Tb and EE for therapeutic purposes in humans.

## Methods

### Animals

All animal care and experimental procedures were approved by the Beth Israel Deaconess Medical Center Institutional Animal Care and Use Committee or the MIT Committee on Animal Care. *Agrp*-IRES-Cre (JAX stock 012899)^37^, Vgat*^lox/lox^* (JAX stock 012897)^37^, NPY-KO (JAX stock 004545)^44^, *Npy*-IRES2-FlpO (JAX stock 030211)^45^, NPY-hrGFP (Jax stock 006417)^46^, *Npy1r*^Cre^ (JAX stock 030544)^47^, *Npy2r*-IRES-Cre (JAX stock 029285) and *Ptger3^Cre^* (Jax stock 035575) mice were maintained on a mixed background.

We used male and female mice aged 10-22 weeks for our experiments. Mice were individually housed in standard plastic cages with standard corn cob bedding with nesting materials on a 12Oh light: 12Oh dark cycle at ambient temperatures ranging between 22 ± 2°C. Mouse chow (Labdiet 5008 irradiated) and water were provided *ad libitum,* unless mice were exposed to food deprivation, as specified in the text.

### Surgery

All surgeries were performed in aseptic conditions. Mice were anesthetized with isoflurane or the mix of ketamine/xylazine (100 and 10 mg/kg, respectively, i.p.) with additional doses of 10% of the initial dose as needed throughout surgery to eliminate the withdrawal reflex. Stereotaxic microinjections into the MnPO (coordinates: AP = +0.43 mm, L = 0.0, DV = −4.6 mm from Bregma) or ARC (coordinates for AAV injection; AP: -1.5 and -1.7 mm, ML: -0.25 and 0 mm, DV: -5.90 mm; coordinates for fiber and cannula implants: AP = −1.6 mm, L = 0.25, DV = −5.85 mm from Bregma). The optical fiber placements in the MnPO were made using the coordinates: AP = +0.43 mm, L = 0.0, DV = −4.58 mm from Bregma.

Mice were implanted with radiotelemetry sensors (TA-F10 or ETA-F10, DSI) in the intraperitoneal space via laparotomy. For the electrocardiogram (HR) recordings, we fixed one of the electrodes attached to the ETA-F10 sensor on the chest muscles and the second electrode in the abdominal region. Then, the muscle layers around the electrodes were closed with absorbable sutures and the abdominal wall was sutured using absorbable sutures. Meloxicam slow-release treatment, for analgesia, was administered prior to surgery. Mice were allowed to recover at least 10 days prior to experimentation. Following recovery, mice showed no signs of discomfort and gained weight normally.

For TeLC injections, mice were anesthetized with isoflurane (1.5-2%) and implanted intra-abdominally with telemetry temperature and activity probes (Unified Information Devices, model UCT-2112). Four craniotomies were performed per animal, followed by four injections (25□nL each) delivered at a rate of 25□nL/sec at AP −1.3□mm and −1.7□mm, ML ±0.35□mm from Bregma, and DV −5.6□mm from the brain surface. Coordinates were adjusted by ±0.1□mm between animals as needed.

### Cloning of Ptger3 Promoter

The Ptger3 promoter sequence was amplified from genomic DNA isolated from C57BL/6J mice, using primers as below:

Primer F (CQO315): gtccaacattaccgccatgttgacactttcacctgtgtgcttacttgg

Primer R (CQO316): atggtggcgaccggtagcatcgaattcCGGGGCCACGGTAGGGCGC

The amplified sequence was digested and ligated into pAAV-GFP/Cre (Addgene, 49056), to replace the CMV promoter and generate pAAV-Ptger3-Promoter-GFP/Cre. Two independent clones were isolated, and the Ptger3 promoter was verified by Sanger sequencing.

### Viral Vectors

For fiber photometry experiments, AAV1-hSyn-DIO-GCaMP6s (Addgene 100845-AAV1; 1.6□×□10^13^ viral genomes□ml^−1^) was delivered into the ARC (100 nL per site above).

For optogenetic experiments, pAAV5-EF1a-DIO-hChR2(H134R)-EYFP-WPRE-hGHpA (AAV5),AAV1-hSyn2-SIO-eOPN3-mScarlet-WPRE (Addgene 125713-AAV1; 1□×□10^13^□viral genomes□ml^−1^) or AAV5-Syn-FLEX-rc[ChrimsonR-tdTomato] (Addgene 62723-AAV5; 2.1□×□10^13^ viral genomes□ml^−1^) was injected bilaterally into the ARC (100 nL per site above).

For TeLC/EYFP injections, pAAV-hSyn-DIO-EGFP and pAAV-hSyn-FLEX-TeLC-P2A-EYFP-WPRE were used. pAAV-hSyn-DIO-EGFP (gift from Bryan Roth, Addgene viral prep # 50457-AAV8) was injected at 2.4×10^^13^ vp/mL. pAAV-hSyn-FLEX-TeLC-P2A-EYFP-WPRE (gift from Fan Wang, Addgene plasmid # 135391 was packaged at Boston Children’s Hospital Core and injected at ∼3.3×10^^12^ vp/mL.

For the AAV-cre_Ptger3 injections in the MnPO, we first packaged the AAV-Ptger3-Promoter-GFP/Cre according to an established protocol^48^. Briefly, five 150-mm dishes were each seeded with 9×10^6^ 293FT cells (Thermo Fisher, R70007) in 20Oml of complete DMEM medium consisting of DMEM (Gibco, 11995-073) supplemented with 10% FBS (Corning, 35-010-CV), 1% MEM Non-Essential Amino Acids (NEAA) (Gibco, 11140-050), 1 mM Sodium Pyruvate (Gibco, 11360070), and 1% GlutaMAX™ Supplement (Gibco, 35050061).

The next day, 40 µg of total plasmid DNA (including pAAV-Ptger3-Prom-GFP/Cre, pUCmini-iCAP-PHP.eB, and pAdDeltaF6) are transfected for each 150-mm dish. At 12-24 h post transfection, the old media was aspirated and replaced with 20 ml of fresh, warmed media containing 5% FBS, 1% NEAA, 1 mM Sodium Pyruvate and 1% GlutaMAX™ Supplement. Cell culture supernatants were collected at 72 h and 120 h post-transfection. Polyethylene glycol (PEG; MW 8,000; Sigma-Aldrich, 89510-1KG-F) was added to the clarified supernatant to a final concentration of 8% (w/v) using the 40% (w/v) PEG stock solution. Samples were incubated overnight at 4O°C with gentle shaking to precipitate viral particles. PEG pellets were collected by centrifugation at 4,000g for 30 min at 4 °C.

Cells were harvested at 120 h post-transfection, and cell pellets were collected by centrifugation at 2,000g for 15 min at room temperature. PEG pellets and cell pellets were treated with SAN nuclease (NEBExpress Salt Active Nuclease; NEB, M0764S) and purified by iodixanol gradient ultracentrifugation using 5%, 25%, 40%, and 60% (w/v) iodixanol solutions (Cosmo Bio USA, AXS-1114542-5). Each gradient consists of 6 ml of 15% iodixanol, 6 ml of 25% iodixanol, 5 ml of 40% iodixanol, and 5 ml of 60% iodixanol prepared in OptiSeal polypropylene tubes (Beckman, 361625). Gradients were sequentially underlaid from low to high density using a 10-ml syringe fitted with an 18-gauge blunt-ended Hamilton needle (Hamilton, 7748-04), taking care to avoid bubble formation. Approximately 6-7 ml SAN-digested lysate was gently layered onto the top of the 15% iodixanol layer using a P1000 pipette. Tubes were filled to the neck with SAN digestion buffer, sealed with black caps and balanced before ultracentrifugation.

Gradients were centrifuged in a Type 70 Ti rotor at 350,000g (58,400 r.p.m.) for 2 h 25 min at 18O°C with slow acceleration (no. 3; the instrument will take 3 min to accelerate to 500 r.p.m., followed by maximum acceleration) and deceleration (no. 9; the instrument will decelerate at maximum speed until it reaches 500 r.p.m., then take 6 min to stop) profiles. Following ultracentrifugation, 4.0-4.5 ml viral fraction was collected from the 40/60% interface and the 40% iodixanol layer using a 5-ml syringe fitted with a 16-gauge needle, while avoiding the protein layer at the 25/40% interface. The collected fraction was diluted into 10 ml DPBS containing 0.001% (v/v) Pluronic F-68 (Gibco, 24040-032) and concentrated using Amicon Ultra centrifugal filter units by centrifugation at 3,000g for 5-8 min at RT until the remaining volume reached 500–1,500 µl Buffer exchange was performed three additional times with 13 ml DPBS containing 0.001% (v/v) Pluronic F-68 for a total of four washes. Final virus preparations (300–500 µl) were transferred to 1.6-ml screw-cap tubes and stored at 4O°C for short-term use or at −80O°C for long-term storage.

For viral genome quantification, unpackaged DNA was digested with DNase I (Sigma, 4716728001), followed by proteinase K digestion (Sigma, 3115887001) to degrade viral capsids and release encapsidated viral genomes. pAAV-Ptger3-Prom-GFP/Cre plasmid DNA was used as the qPCR standard. Three independent sets of eight 1:10 serial dilutions were prepared, corresponding to 5-10 × 10^^8^ to 5-10 × 10^^1^ single-stranded DNA molecules/µl. qPCR reactions contained 12.5 µl SYBR Green master mix, 9.5 µl UltraPure water, 0.5 µl forward primer, 0.5 µl reverse primer and 2 µl template DNA. Reactions were performed using the following cycling conditions: 95O°C for 10 min, followed by 40 cycles of 95O°C for 15 s and 60O°C for 40 s. The final viral genome titers were AAV 3.67 ×10^^12^ gc/ml.

A mix of AAV-cre_Ptger3 and the AAV8-hSyn-DIO-hM3D(Gq)-mCherry (AAV-hM3Dq) produced at the University of North Carolina virus core was used in volumes of ∼100 and ∼300 nL^49^.

pAAV5-DIO-taCasp3-TEVp (AAV-Casp) was packaged and obtained by Addgene. It was delivered into the MnPO of each mouse by a single brain injection of a 1:1 mix of AAV-cre_Ptger3 and AAV-Casp; we also added the non-cre-dependent AAV-GFP at a 1:2 ratio to label the injection sites.

### Brain injections of NPY

For the MnPO injections of NPY, we used a Hamilton syringe to deliver 4 μg in 1 μL of NPY (Tocris Catalog #1153) or 50% DMSO (control). We performed a single injection of NPY after the mice were restricted from food access for 2 hours. We returned the food 3 hours after delivering NPY in the MnPO of these mice.

### Body temperature, locomotor activity and heart rate recordings

Tb, LMA and HR was recorded using the radiotelemetry DSI system. The signal was sent from the telemetry sensors previously implanted in mice to DSI receivers and converted using the PhysioTel HD and PhysioTel (DSI) hardware. We used the mean of core temperature, locomotor activity, and heart rate every 1-5 min.

For TeLC experiments, telemetry receiver platforms (UID Mouse Matrix) were used for continuous temperature and activity monitoring. Raw data were exported from the MouseMatrix platform Software (Unified Information Devices, MouseMatrix 2.153.1.0) and the median value calculated per minute. All telemetry data were processed using a custom R package developed for UID data parsing and analysis, available at https://github.com/matiasandina/uid

### Food Deprivation

Mice were kept at 22 ± 2 °C and food deprived for 24-48h, starting at (ZT6) in some of our fiber photometry tests or the beginning of the dark phase (ZT12), as indicated in the text. Water was provided *ad libitum*.

### Energy expenditure measurements

We used the Promethion metabolic cages by Sable Systems International to measure oxygen consumption (VO2), carbon dioxide production (VCO_2_), and respiratory exchange ratio (RER) every 3-5 minutes in a temperature-controlled cabinet 23 +/- 0.2 °C.

### Fiber photometry

Most fiber photometry recordings were performed as previously described^50^. Mice were singly housed in optogenetics-compatible cages (BioDAQ, Research Diets) and fiber photometry was performed using one color time-division multiplexed illumination using the pyPhotometry acquisition system^51^. 465 nm (CLED_465 nm; Doric Lenses) and 405 nm (CLED_405 nm; Doric Lenses) LEDs were used as excitation sources for GCaMP6 and isosbestic signals respectively.

Excitation light was transmitted onto the sample via an optic fiber cable (1 m long; 400 μm diameter NA 0.48; Doric Lenses) coupled to the implanted optic fiber. Emission light was collected via the same optic fiber cable and separated from excitation light using a fluorescence mini-cube (FMC4_IE(400_410)_E(460_490)_F(500_550)_S; Doric Lenses). Recordings were performed at 20Hz and the signal was collected by the photodetector integrated into the mini-cube.

Other experiments were performed using the same LEDs and Doric mini-cube but, signals were acquired using the Spike2 software. Post-hoc analysis of the acquired data was also performed using Spike2 analysis that involved low pass filtering of acquired signal and normalization of the data to the baseline or other control recordings^17^.

For the analysis of all fiber photometry data, data were analyzed using custom Python (Python 3.12) scripts. All data were median-filtered and low-pass-filtered (10Hz cutoff). Motion correction was performed by finding a linear best fit to the 405 nm signal and subtracting this from the 465 nm signal. ΔF/F was calculated as (465 nm – 405 nm fitted signal) / F0, where F0 is the low-pass filtered 465 nm signal (0.001 Hz cutoff) of the whole recording session. Traces were smoothed using a 30-minute running average. We defined torpor entry as 30 minutes before the body temperature went below 33 °C and, baseline as 30 minutes before the torpor entry. Z score was calculated across the whole trace and then baseline-corrected by subtracting the mean of the baseline period from the whole trace.

### Photostimulation and Photoinhibition

Photostimulation and Photoinhibition were delivered at 4–12 mV (4-8 mV for the eOPN studies and 8-12 mV for the ChR2 and Chrimson photostimulation) by using a blue light laser (470 nm; LaserGlow), and the patterns of stimulation (10-20Hz and the time of each stimulation) were controlled by Spike2 8.01 software. For optogenetically activating the ARC^AgRP^→MnPO circuit, we used 20 Hz, 2-sec laser on and 3-sec off continuously. For photostimulation of the POA cell bodies, we used 10 Hz, 2-sec laser on and 3-sec off continuously. For photoinhibition of the ARC^AgRP^→MnPO pathway, we used 10 Hz, 2-sec laser on and 3-sec off continuously.

### Thermal imaging

Thermal imaging was performed during the light phase using an infrared camera (FLIR E4). Before the recordings, the back, neck and head of the mouse was shaved under low dose of ketamine/xylazine. Following at least 2 days of recovering from anesthesia and 24 hours for habituation, we recorded videos to analyze the changes in skin temperature corresponding to BAT (TBAT) and tail (Ttail) temperatures. TBAT was determined by using a mean of the temperature between the right and left scapula) and Ttail by using tree different points from the body to the most distal region of the tail. We used FLIR tools software for all analysis.

### Anatomical tracers

As previously described^19,52^, cholera toxin subunit B (CTb; 1–2% diluted in saline; catalog #103A, List Biological Laboratories) was used as retrograde anatomical experiments. We injected a volume of 6–15 nl into the MnPO of mice.

AAV8-CMV-FLEX-TVAmCherry-2A-oG and EnvA pseudotyped rabies vectors were obtained by Salk Viral Vector core, and both vectors were injected at ∼100 nL per MnPO injection.

### Perfusion and brain sectioning

After performing the experimental protocols, mice were deeply anesthetized with chloral hydrate (1.5% BW i.p. 7% solution) and transcardially perfused with 30ml phosphate buffer saline and then 30ml 10% buffered formalin (Fisher Scientific; Catalog # SF100-20). Brains were extracted and post fixed overnight in 10% formalin and then stored in 20 or 30% sucrose until sectioned using a freezing microtome (20 or 40 µm coronal sections into 3 series) to perform immunohistochemistry or RNAscope insitu hybridization. Following sectioning, tissue was stored at 4°C in PBS containing the preservative sodium azide or at -80°C mounted on the slides until processed for histology.

For TeLC experiments, animals were deeply anesthetized and perfused transcardially with 4% paraformaldehyde (PFA). Brains were extracted and post-fixed overnight at 4O°C in 4% PFA. Coronal sections (50Oµm) were cut using a vibratome (Leica VT1200S), and sections encompassing the arcuate nucleus were selected for immunohistochemistry.

### Injection sites

In order to confirm the injection sites, one series of the tissue was mounted and cover slipped with hard-set mounting media (Vectashield) for native fluorescence (GFP or mCherry). Only animals with the histological confirmation of the region targeted were used for analyses of experimental groups.

### Immunohistochemistry (IHC)

We used a free-floating method to label CTb and cFos in the brain sections^19,52^. In brief, the tissue was rinsed and incubated in a blocking solution of 10% horse serum. Then, we incubated the tissue with the primary antibody for CTb (1:10,000, list biologicals, catalog #703)) or Fos (AB5, 1:10,000; Oncogene Sciences, catalog #484) overnight. The following day, we used the appropriate secondary antibodies, rinsed the tissue, and mounted it onto glass slides. The slides were dried and coverslipped with Vectashield before imaging.

For TeLC-EYFP and EGFP staining, arcuate sections were blocked for 45Omin at room temperature in 10% normal donkey serum (NDS; Sigma-Aldrich, D9663) diluted in 0.4% PBS-Triton. Primary antibody incubation was performed overnight at 4O°C with agitation using chicken anti-GFP (Aves Labs, GFP-1020; 1:1000) in 2% NDS. Following three 10-min washes in 0.4% PBS-Triton, sections were incubated with Alexa Fluor 488-conjugated donkey anti-chicken secondary antibody (ThermoFisher, A78948; 1:500 in 2% NDS) for 2Oh at room temperature. Following three 10-min washes in 0.4% PBS-Triton, sections were floated onto glass slides and coverslipped. All blocking and antibody solutions were prepared in 0.4% PBS-Triton.

### ISH

In situ hybridization assays were performed using RNAscope Multiplex Fluorescent Reagent Kit v2 (Catalog#323100,ACD,CA) following the protocol, which we have established as a reliable method in our laboratory using rodent brains^17,19^. One series of brain sections from animals used for monosynaptic tracing (EnvA pseudotyped rabies) was used to label the mRNA for Npy1r (neuropeptide Y receptor 1) using RNAscope Probe Mm-Npy1r-C2 (Catalog#427021); Agrp (Agouti gene related protein) using RNAscope Probe Mm-AGRP-C2 (Catalog# 400711-C2) and Kiss1 (KiSS-1 metastasis-suppressor) using RNAscope Probe Mm-Kiss1-C1 (Catalog # 500141) in respective experiments

### Histological imaging

We used V200 Olympus slide scanner and Olyvia viewer software V4.1 for imaging the brain tissue from animals injected with AAVs and anatomical tracers. Only animals with the histological confirmation of the region targeted were used for analyses. For confocal microscopy, we used Leica LAS X Confocal and the Leica Microscope Platform. For TeLC experiments, sections were scanned using a TissueFAX SL slide scanner.

### Statistical analysis

Data are presented as mean ± SEM. The statistical test used, statistical significance and number of animal subjects per group are reported in the results and figure legends. The data were considered to be statistically significant when p<0.05. Unless otherwise noted, statistical analyses were performed using GraphPad Prism (GraphPad Software). Following a test for normality (D’Agostino and Pearson omnibus test), significant differences were determined using either repeated measures of one-way ANOVA or two-way ANOVA, both followed by Bonferroni’s correction. For statistical tests between two groups, a two-sided t-test was used. For TeLC experiments, non-parametric Wilcoxon rank-sum tests were used to test two-group samples. Fisher’s Exact Test was used to test likelihood given a 2×2 contingency table. A mixed effects linear model was fitted to the minimum temperature shown on each session followed by post-hoc contrasts. In this case, p values Holm-corrected within contrast family. Statistical analysis was performed in R using custom scripts.

## Supporting information

Ext data Fig 1

Ext data Fig 2

Ext data Fig 3

Ext data Fig 4

Ext data Fig 5

xExt data Fig 6

Ext data Fig 7

Ext data Fig 8

Ext data Fig 9

Ext data Fig 10

Ext data Fig 11

Ext data Fig 12

## Data availability

The data that support the findings in this study are available from the corresponding author upon request.

## Acknowledgments

We thank Dr. Jon Resch and Carrie Mahoney for their insights and for sharing resources. We also thank John Kim, Quan Ha, Marissa Cortopassi, Keilana Godhwani, Cecilia Villareal, Perla Chacon, and Ashlin Kelly for their excellent technical assistance.

Funded by BIDMC The Mathers Foundation # MF-2204-02555 (Machado) and Nutrition Obesity Research Center at Harvard # NIDDK-P30-DK040561 (Machado). Dr. Costa received support from the São Paulo Research Foundation (FAPESP). This work was supported by the NIH (P30DK046200, P30DK135043, R01DK122976, R01DK075632, R01DK089044, R01DK111401, R01DK134427, and R01DK096010 to Lowell, DP2DK136123-02 to Hrvatin, and R00AG076739 to Qiu). The Charles A. King Trust Postdoctoral Research Fellowship Program (Douglass), a BBRF Young Investigator Grant (Kucukdereli), and Alzheimer’s Association (Qiu).

## Author contributions

AMD, CBS, BBL and NLSM contributed to the concept and design of the study. AMD, NL, DM, LHAC, OR and NLSM performed the experiments. AMD, NL, HK, DM, LHAC, ORP, ASB, YY, CQ, SH, CBS and NLSM performed data and/or histological analysis. SH and MA conceptualized, performed, and analyzed the experiments using TeLC. SH and MA also analyzed the transcriptomic dataset from the Allen Brain Institute^31^. CQ and YY cloned the Ptger3 promoter and packaged the AAV-Ptger3-Promoter-GFP/Cre. SSB performed the ISH. AMD, CBS, BBL and NLSM wrote the manuscript. All authors edited and approved the final version.

## Legends Extended Data

**Ext data Fig 1: The activity of ARC^AgRP^ neurons is inversely related to body temperature.**

(A) Individual changes of calcium signaling and body temperature in 3 recorded animals across fed AL, fasted, and refed conditions.

**Ext data Fig 2: ARC^AgRP^ neurons are required for fasting-induced torpor**.

(A) Mean body temperature of control (ARC^AgRP-EGFP^) and ARC^AgRP-TeLC^ groups during the baseline recording (EGFP: 36.92 ± 0.08 °C; TeLC: 36.87 ± 0.09 °C; Welch’s t-test, tO17.4O = 0.41, p = 0.690).

(B) Individual body temperature traces for each animal in control (EGFP) and TeLC groups during the fasting session.

(C) Total time in torpor during the fasting session (EGFP: 302.67 ± 63.92 min; TeLC: 75.44 ± 52.27 min; Wilcoxon’s rank sum test, p = 0.013).

(D) Weight of mice (EGFP, gray; TeLC green) before fasting session (EGFP: 21.31 ± 0.40 g; TeLC: 24.57 ± 0.73 g; Welch two sample t-test, p = 0.007).

(E) Linear regression on minimal body temperature during the fasted session. Single additive factors include vector and group centered weight. After centering by group weight, weight itself was not a significant predictor of minimal temperature during fasting (β = 0.12 ± 0.29, p = 0.696).

(F) Baseline mean body temperature for B6/C57 EGFP and B6/C57 TeLC mice was not significantly different (EGFP: 37.24 ± 0.11 °C; TeLC: 37.05 ± 0.07 °C; Welch’s t-test, tO11.3O = 1.46, p = 0.171)

(G) Mean body temperature recorded from B6/C57 TeLC (green, n=10) or B6/C57 EGFP (gray, n=7) mice during AL or FD conditions.

(H) Percentage of B6/C57 mice entering long-duration torpor (EGFP: 3/7; TeLC: 6/10; Fisher’s Exact test, odds ratio = 0.52, p = 0.883)

(I) Minimum body temperature during baseline and fasting sessions for B6/C57 EGFP and B6/C57 TeLC mice. There was no vector-by-session interaction (β = 0.43 ± 0.70°C, tO15O = 0.62, p = 0.546). Across vectors, fasting reduced minimum body temperature (β = −4.77 ± 0.34 °C, tO16O = −14.13, p < 0.001), while we found no significant differences in EGFP and TeLC groups overall (β = −0.18 ± 0.47 °C, tO15O = −0.38, p = 0.711).

(J) Mean torpor bout duration during the fasting session for B6/C57 EGFP and B6/C57 TeLC mice (EGFP: 75.79 ± 21.35 min; TeLC: 110.10 ± 28.23 min; Wilcoxon’s rank sum test, p = 0.826).

In panels B, C, and D, each dot is a single animal. Data are represented as Mean ± SEM.

**Ext data Fig 3: ARC^AgRP^ synaptic terminals in the MnPO show increased calcium signaling during fasting-induced torpor.**

(A) Individual changes of calcium signaling and body temperature in 3 recorded animals across 24-48 hours of food deprivation.

**Ext data Fig 4: Optogenetic activation of the ARC^AgRP^**→**MnPO pathway causes an increase in food intake in a subset of mice.**

(A) Schematic of approach using ChR2-mediated optogenetic activation of the ARC^AgRP^ — MnPO circuit in mice.

(B) Quantification of feeding in mice subjected to 4-h activation of the ARC^AgRP^ — MnPO (STIM) or controls (SHAM): 120 ± 48 mg SHAM AL vs. 385 ± 152 mg STIM AL, unpaired t-test, p=0.067, n=5.

(C) Representative images of the injection sites in the ARC and the fiber placement in the MnPO.

Data are represented as Mean ± SEM.

**Ext data Fig 5: Fasting-induced torpor in *Agrp*-IRES-cre, *Agrp*-IRES-cre;NPYKO and *Agrp*-IRES-cre;Vgat^fl/fl^ mice.**

(A) Schematic of approach using eOPN3 for inhibition of the ARC^AgRP^ — MnPO pathway in mice (Top panel).

(B) Changes in the locomotor activity of mice in response to fasting during SHAM condition or inhibition of the ARC^AgRP^ — MnPO pathway (left). The mean locomotor activity of mice under SHAM and STIM conditions during ZT22 to ZT6 (when the propensity for torpor bouts is high) does not show statistically significant differences (2.0 ± 1.2 SHAM FD vs. 3.33 ± 1.9 STIM FD, paired t-test p=0.15).

(C) Representative images of AAV-eOPN3 expression (red) in AgRP-expressing neurons in the arcuate nucleus (bottom left) and the fiber placement lateral to the synaaptic terminals in the MnPO (bottom right).

(D) During food deprivation, control mice show similar changes in Tb during SHAM or optogenetic stimulation (bottom panel).

(E) Illustration of mice subjected to a 24-h period of food deprivation, while their Tb were monitored by a telemetry system.

(F) Quantification of the average number torpor bouts (1.3 ± 0.2 bouts *AgRP*-IRES-cre vs 0.75 ± 0.5 bouts *AgRP*-cre;NPYKO vs 1.8 ± 0.6 bouts *AgRP*-cre; Vgat^fl/fl^, n=6, 4 and 5 respectively, One-way ANOVA, followed by Bonferroni’s post hoc test, p> 0.05). Duration of torpor bouts (165 ± 25 min *AgRP*-IRES-cre vs 65 ± 44 min *AgRP*-cre;NPYKO vs 354 ± 102 min *AgRP*-cre; Vgat^fl/fl^, n=6, 4 and 5 respectively, One-way ANOVA, followed by Bonferroni’s post hoc test, p> 0.05) of torpor bouts displayed by *Agrp*-IRES-cre, *Agrp*-IRES-cre;Vgat^fl/fl^ mice, and *Agrp*-IRES-cre;NPYKO mice during a 24-h period of food deprivation.

Data are represented as Mean ± SEM.

**Ext data Fig 6: *Agrp*-IRES-cre;Vgat^fl/fl^ mice show hypothermic response induced by activation of the ARC^AgRP^→MnPO pathway.**

(A) Schematic of approach using ChR2-mediated optogenetic activation of the ARC^AgRP^— MnPO circuit in *AgRP*-cre;Vgatfl/fl mice (top panel). Representative images of ChR2 expression (red) in AgRP-expressing neurons in the arcuate nucleus (bottom left) and their synaptic terminals in the MnPO below the area of the fiber placement (bottom right).

(B) Changes in body temperature recorded from the same *AgRP*-cre;Vgatfl/fl mice during activation of the ARC^AgRP^ — MnPO (STIM) or controls (SHAM) in mice fed ad libitum (AL).

(C) Quantification of the changes in mean Tb after activation of the ARC^AgRP^ — MnPO (STIM) or controls (SHAM) in mice fed ad libitum, 1-h STIM: 37.16 ± 0.03 °C SHAM vs 35.45 ± 0.1 °C STIM, paired t-test, p=0.0005; 2-h STIM 36.87 ± 0.02 °C SHAM vs 32.04 ± 0.28 °C STIM, paired t-test, p=0.0005; 3-h STIM 36.66 ± 0.15 °C SHAM vs 30.52 ± 0.65 °C STIM, paired t-test, p=0.0025; 4-h STIM 36.78 ± 0.07 °C SHAM vs 31.39 ± 0.31 °C STIM, paired t-test, p=0.0004; n=4. Data are represented as Mean ± SEM.

(D) Representative images of the injection sites in the ARC and the fiber placement in the MnPO.

**Ext data Fig 7: NPY and GABA are neuromodulators of the hypothermic response induced by activation of the ARC^AgRP^**→**MnPO pathway**

(A) Schematic approach to deliver NPY in the MnPO, while recording Tb changes in WT mice (left), and a representative image of the cannula placement and injection site in the MnPO (right).

(B) Mean body temperature recorded after one single injection of NPY or vehicle (control) in the MnPO of mice (Mean Tb 1-h after local MnPO injection: 37.68 ± 0.2 °C SEM vehicle vs 33.07 ± 0.7 °C SEM NPY, two-way ANOVA, followed by Bonferroni’s post hoc test, *p<0.0001, n=5).

(C) Schematic of approach using ChR2-mediated optogenetic activation of the ARC^AgRP^ — MnPO circuit in *AgRP*-cre;NPYKO mice

(D) Changes in body temperature recorded from the same *AgRP*-cre;NPYKO mice during activation of the ARC^AgRP^ — MnPO (STIM) or controls (SHAM) in mice fed ad libitum (AL).

(E) Quantification of the changes in mean Tb after activation of the ARC^AgRP^ — MnPO (STIM) or controls (SHAM) in mice fed ad libitum. Mean Tb 4-h STIM: 36.51 ± 0.4 °C SHAM vs 35.52 ± 0.1 °C STIM, paired t-test, p=0.04, n= 2 female and 1 male.

(F) Representative image of the AAV-ChR2 expression in the ARC (top left) and fiber placement in the MnPO (top right). Illustrations of the injection sites in the ARC (bottom left) and the fiber placement in the MnPO (bottom right) of *AgRP*-cre;NPYKO mice.

(G) Schematic approach using mCherry as control for optogenetic activation of the ARC^AgRP^ — MnPO circuit in *AgRP*-cre;NPYKO;Vgatfl/fl mice mice.

(H) Change in body temperature recorded from *AgRP*-cre;NPYKO;Vgatfl/fl mice injected with ChR2 or mCherry (control) in ARC^AgRP^.

(I) Stimulation of the ARC^AgRP^→MnPO pathway in mice expressing ChR2 (n=5) did not alter body temperature when compared to controls expressing mCherry (n=3), two-way ANOVA, followed by Bonferroni’s post hoc test, F_(3,24)_=1.325, p=0.28, n=5.

(J) Representative image of the AAV-mCherry injection site in the ARC (top left) and fiber placement in the MnPO (top right). Illustrations of the injection sites in the ARC (bottom left) of *AgRP*-cre;NPYKO;Vgatfl/fl mice and the fiber placement in the MnPO (bottom right).

Data are represented as Mean ± SEM.

**Ext data Fig 8: The hypothermic response caused by activating the ARC^AgRP^**→**MnPO pathway is attenuated in mice lacking MnPO^EP3R^ neurons.**

(A) Schematic figure of an AAV construct enabling the expression of Cre recombinase in neurons that express the *Ptger3* gene, AAV-cre_Ptger3.

(B) After the injection of an AAV mix containing AAV-cre_Ptger3 and AAV-DIO-hM3Dq in the MnPO of wild-type mice (n=3), ISH was used to validate AAV-hM3Dq expression in MnPO^EP3R^ neurons. Left image, overview of the POA, followed by higher magnification images of example cells that co-express hM3Dq and *Ptger3* mRNA. Scale left = 1mm. Scale right = 500um.

(C) Schematic of approach using an AAV mix containing AAV-cre_Ptger3 and AAV-DIO-hM3Dq to activate MnPO^EP3R^ neurons in wild-type mice.

(D) Individual Tb traces showing the ability of wild-type mice injected with an AAV mix containing AAV-cre_Ptger3 and AAV-DIO-hM3Dq in the MnPO to produce hypothermia in response to CNO treatment (0.3 mg/kg, n=5). The hypothermic response induced by this approach causes profound hypothermia, which is also observed in Ptger3^Cre^ mice injected with AAV-DIO-hM3Dq in the MnPO and treated with CNO (0.3 mg/kg, n=2).

(E) Schematic of approach using ChR2-mediated optogenetic activation of the ARC^AgRP^ — MnPO circuit in mice after ablation of MnPO^EP3R^ neurons.

(F) Representative images of the injection site in the ARC and MnPO, as well as the fiber placement in the MnPO. Scale bar = 100um.

(G) Mean body temperature recorded from *AgRP*-cre mice during ChR2-mediated activation of the ARC^AgRP^ →MnPO pathway for 4-h (ZT3-ZT7) caused hypothermic responses in animals fed *ad libitum* (AL, n=5). The hypothermic response caused by the optogenetic activation of the ARC^AgRP^ →MnPO pathway was attenuated in *AgRP*-cre mice injected with an AAV mix containing AAV-cre_Ptger3 and AAV-Casp9 in the MnPO.

(H) Quantification of the hypothermic responses of mice subjected to activation of the ARC^AgRP^→MnPO (Mean Tb change on an hourly basis), 2h: -1.46 ± 0.13 °C STIM AL vs -0.91 ± 0.08 °C STIM AL ablation MnPO^EP3R^, unpaired t-test, p=0.008 and 4h: - 2.16 ± 0.25 °C STIM AL vs -1.47 ± 0.11 °C STIM AL ablation MnPO^EP3R^, unpaired t-test, p=0.01.

Data are represented as Mean ± SEM.

**Ext data Fig 9: Analysis of MERFISH dataset centered in neuropeptide receptors in POA.**

(A-C) MERFISH expression data showing Npy1r (A), Npy2r (B), and Mc4r (C). Each dot represents a cell in the MERFISH dataset. For illustration purposes, expression counts were only mapped onto a colored scale for cells located inside the POA and neighboring regions.

(D) UMAP plot of all cells in the POA. Cells identified as neurons are highlighted in pink.

(E) UMAP plots of neurons selected from (D). Panels show UMAP plots for Slc17a6 and Slc32a1 expression mapped onto the color scale, as well as the Leiden clustering with annotated clusters. The cluster name acronym comprises neuronal class (E, excitatory; I, inhibitory), followed by cluster number.

(F) Dotplot showing top 3 identifying genes for each cluster found using the cosine similarity-based method for marker gene identification. The marginal barplot shows the cluster sizes. The fraction of cells identified in each cluster expressing each gene is mapped to the size of circles, and the mean expression normalized by gene is mapped to the color scale.

(G) Gene expression for each one of the receptors of interest in each neuronal population.

**Ext data Fig 10: Monosynaptic rabies tracing of inputs to MnPOEP3R neurons.**

(A) Schematic of the tracing strategy: Ptger3-IRES-Cre mice received Cre-dependent helper AAVs (TVAmCherry-2A-oG) in the MnPO, followed by EnvA-pseudotyped, glycoprotein-deleted rabies virus (SADΔG-EGFP) to label direct monosynaptic inputs onto MnPO^EP3R^ neurons.

(B) Representative images of starter cells in the MnPO and EGFP-labeled presynaptic neurons in the preoptic region (POA), periventricular nucleus of the hypothalamus (PVH), suprachiasmatic nucleus (SCN), subparaventricular zone (SPZ), arcuate nucleus (ARC), dorsomedial hypothalamus (DMH), and premammillary nucleus (PMV). 3V, third ventricle.

(C) Semi-quantitative distribution of rabies-labeled input neurons across the brain in four mice. None, no labeled neurons; +, sparse (<5 neurons); ++, moderate (5–20 neurons); +++, dense (>20 neurons).

(D) RNAscope in situ hybridization for *Agrp* and *Kiss1* mRNA combined with EGFP immunostaining in the arcuate nucleus. Rabies-labeled (EGFP+) neurons do not colocalize with *Agrp* mRNA+ cells, indicating that ARC^AgRP^ neurons do not provide direct monosynaptic input to MnPO^EP3R^ neurons. Scale bar = 25um.

**Ext data Fig 11: Optogenetic activation of the NPY2R-expressing POA neurons prevents fasting-induced torpor**

(A) Schematic of approach using ChR2-mediated optogenetic activation of the POA^Npy1r^ neurons (top) and individual Tb recording of male (blue) and female (magenta) mice during SHAM or STIM condition in food deprived (FD) mice (bottom).

(B) Schematic of approach using blue-light activation of the MnPO (top) as control. Individual Tb recording of 2 female (magenta) mice during SHAM or STIM condition in food-deprived (FD) mice is shown (bottom).

(C) Schematic of approach using ChR2-mediated optogenetic activation of the POA^Npy2r^ neurons (top) and a representative image of the injection site and fiber placement in the POA (bottom). Scale bar = 100um.

(D) During the FD condition, all *Npy2r*-expressing mice (4 male mice) showed at least one torpor bout in SHAM condition, while activating the POA^Npy2r^ neurons in these mice precluded fasting-induced torpor in 2 mice (on the left, the individual traces are shown). Evaluation of group data (right) before and after STIM in the FD condition did not show a statistically significant change in mean minimal Tb: 29.74 ± 1.11 SHAM FD vs. 32.78 ± 0.63 °C STIM FD, paired t-test, p=0.1, n=4).

(E) During the FD condition, all *Npy2r*-expressing mice (2 female mice) showed at least two torpor bout in SHAM condition, while activating the POA^Npy2r^ neurons in these mice precluded fasting-induced in both mice (on the left, the individual traces are shown). Evaluation of group data (right) before and after STIM in the FD condition did not show a statistically significant change in mean minimal Tb: 29.61 ± 0.11 SHAM FD vs. 35.38 ± 0.1 °C STIM FD, n=2).

Data are represented as Mean ± SEM.

**Ext data Fig 12: Schematic representation of a central pathway required for fasting-induced torpor.** We propose that in response to low energy stores, activation of ARC^AgRP^ neurons causes the simultaneous release of NPY and GABA in the MnPO, which in turn inhibits GABAergic interneurons that express the NPY1 and NPY2 receptors. In this model, the inputs from ARC^AgRP^ neurons in the MnPO region result in the disinhibition of the MnPO^EP3R^ neurons, leading to an indirect inhibition of the thermogenic neurons in the DMH/DHA, allowing torpor generation.

