## Supplementary figures and images for "A Hypothalamic Pathway for Energy Conservation"

### Ext data Fig 1

#### AgRP neuron Somas

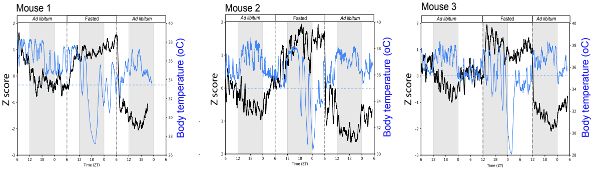


**ARC$^{AgRP}$ neuron activity**
**Body Temperature**

### Ext data Fig 2

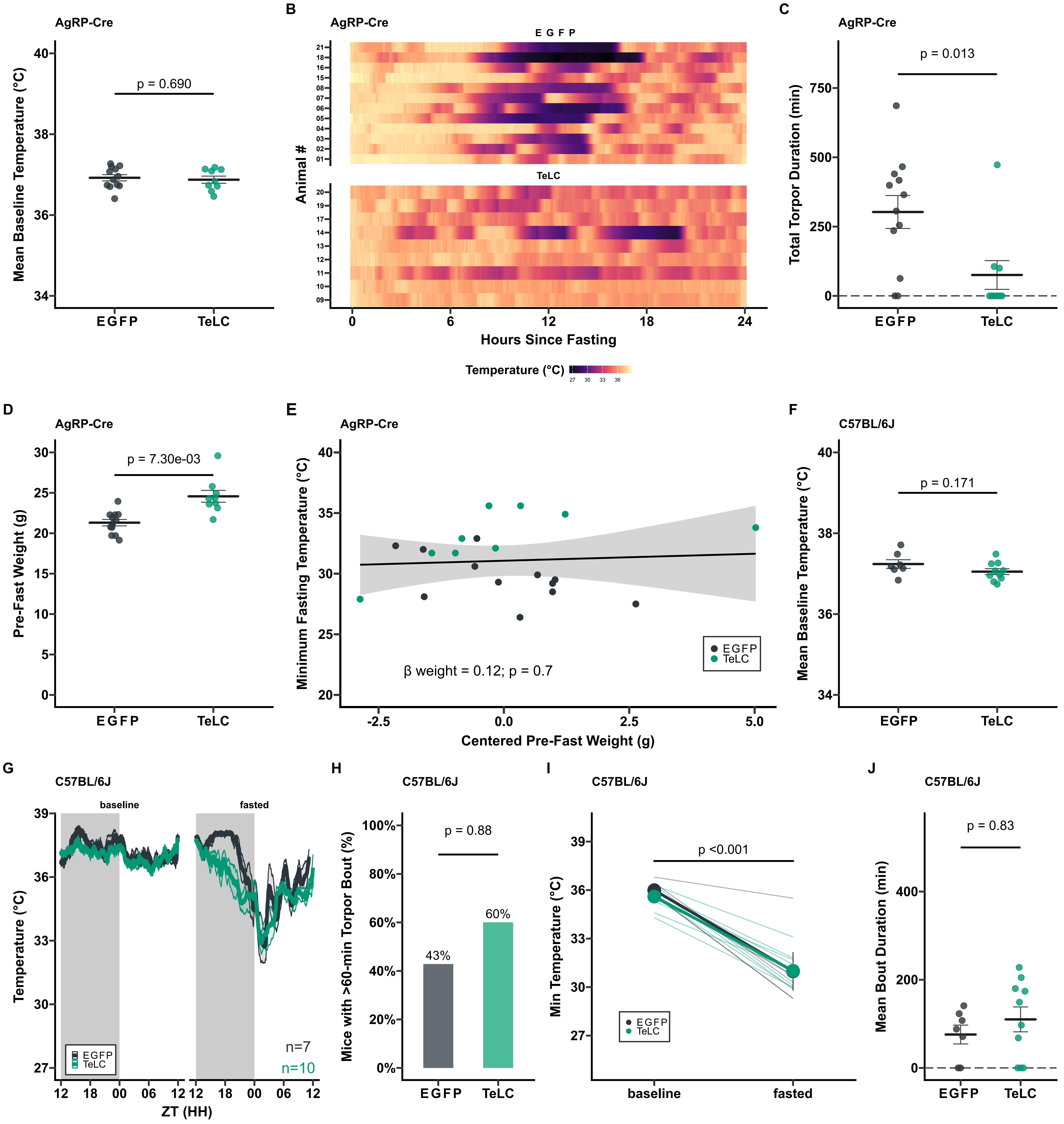

### Ext data Fig 3

Extended Data Fig. 3

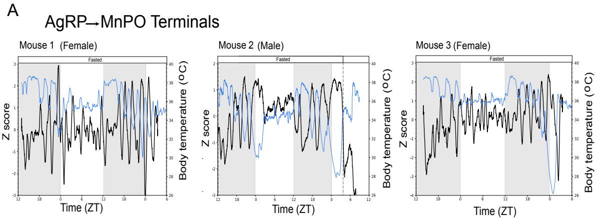

### Ext data Fig 4

Extended Data Fig. 4

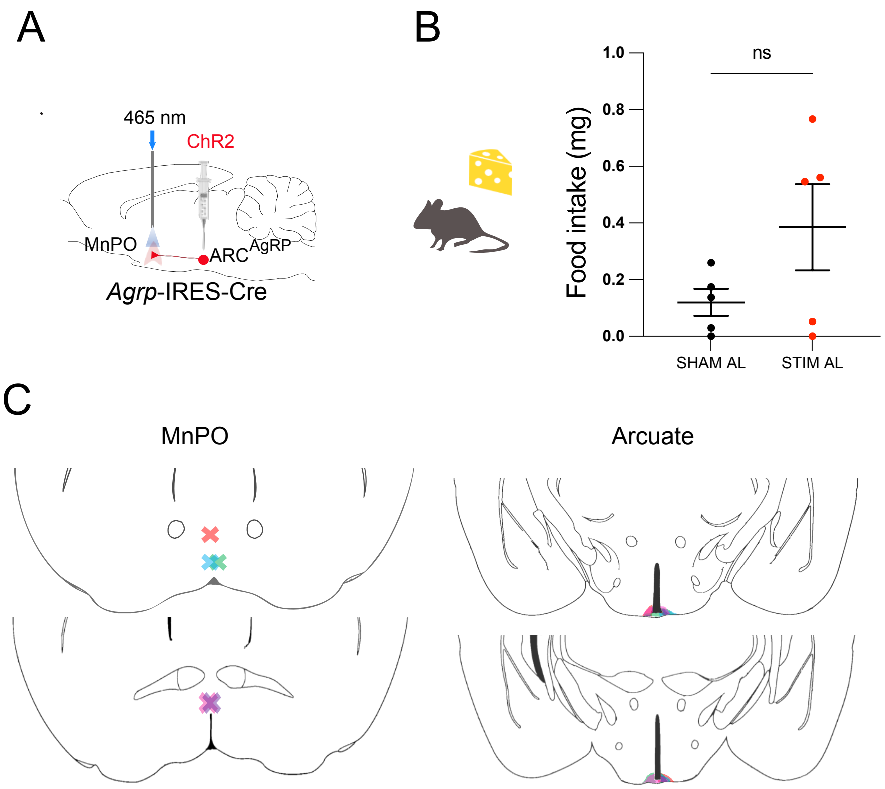

### Ext data Fig 5

#### Extended Data Fig. 5

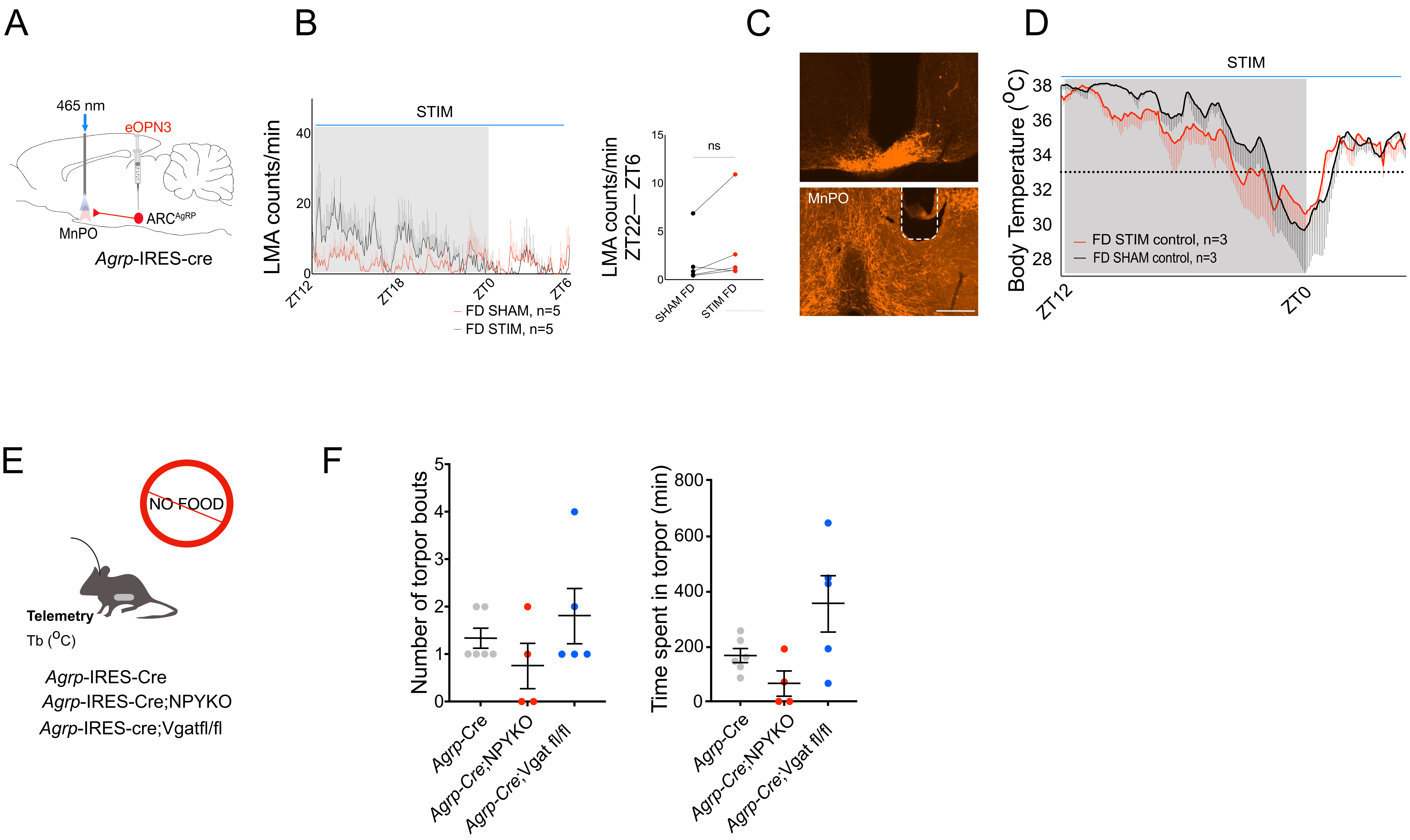

### Ext data Fig 7

#### Extended Data Fig. 7

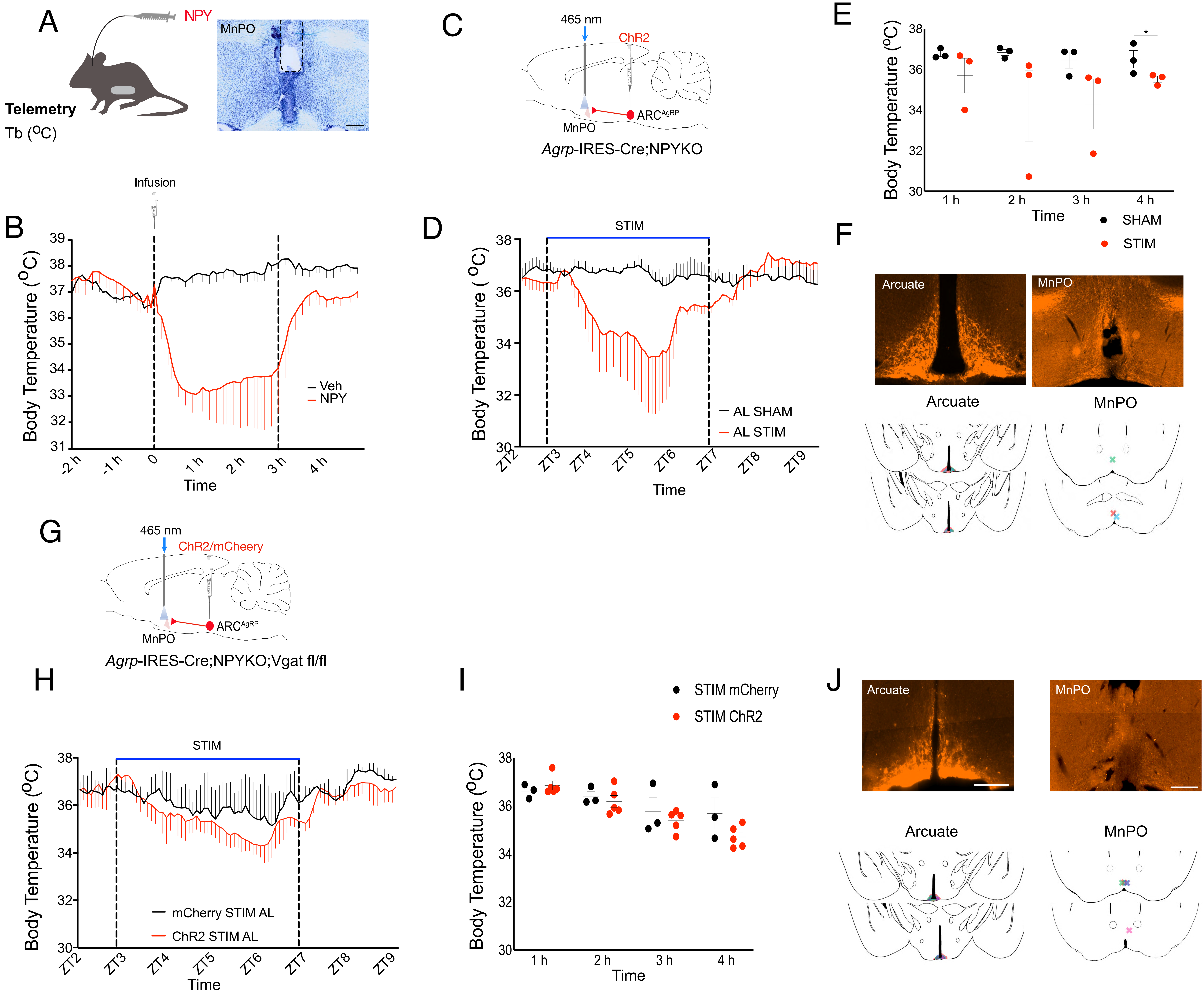

### Ext data Fig 8

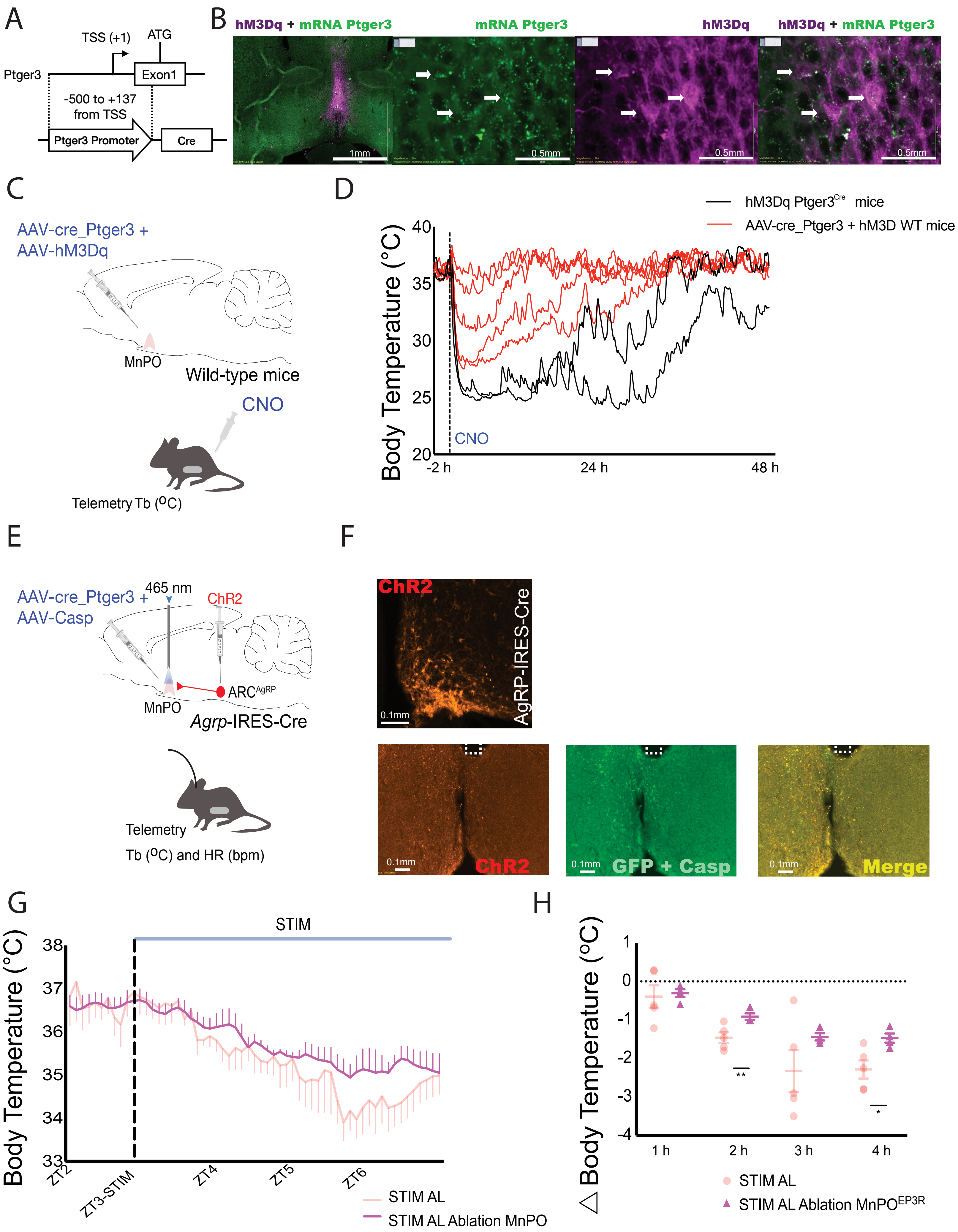

### Ext data Fig 9

Ext data Fig 9

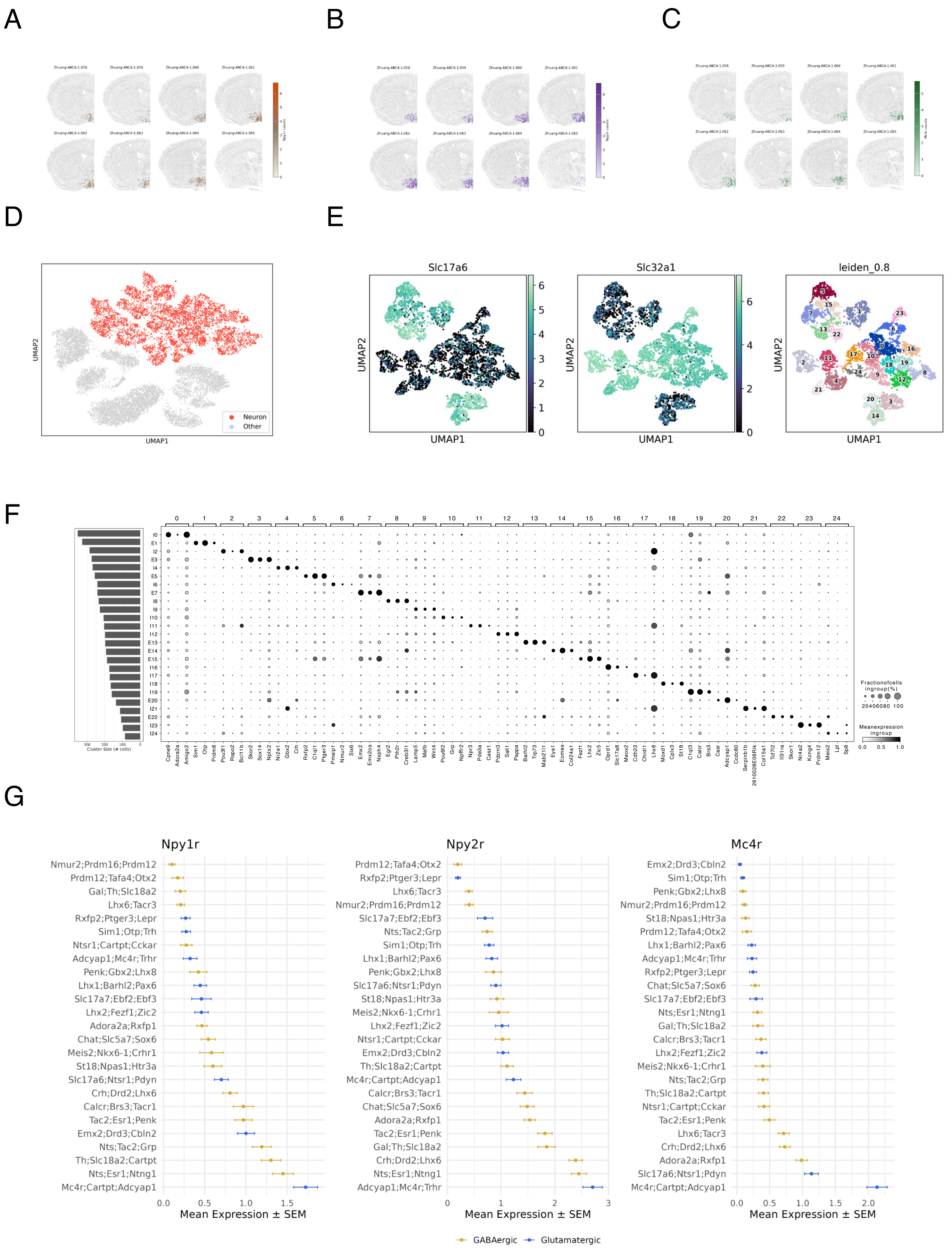

### Ext data Fig 11

Extended Data Fig. 11

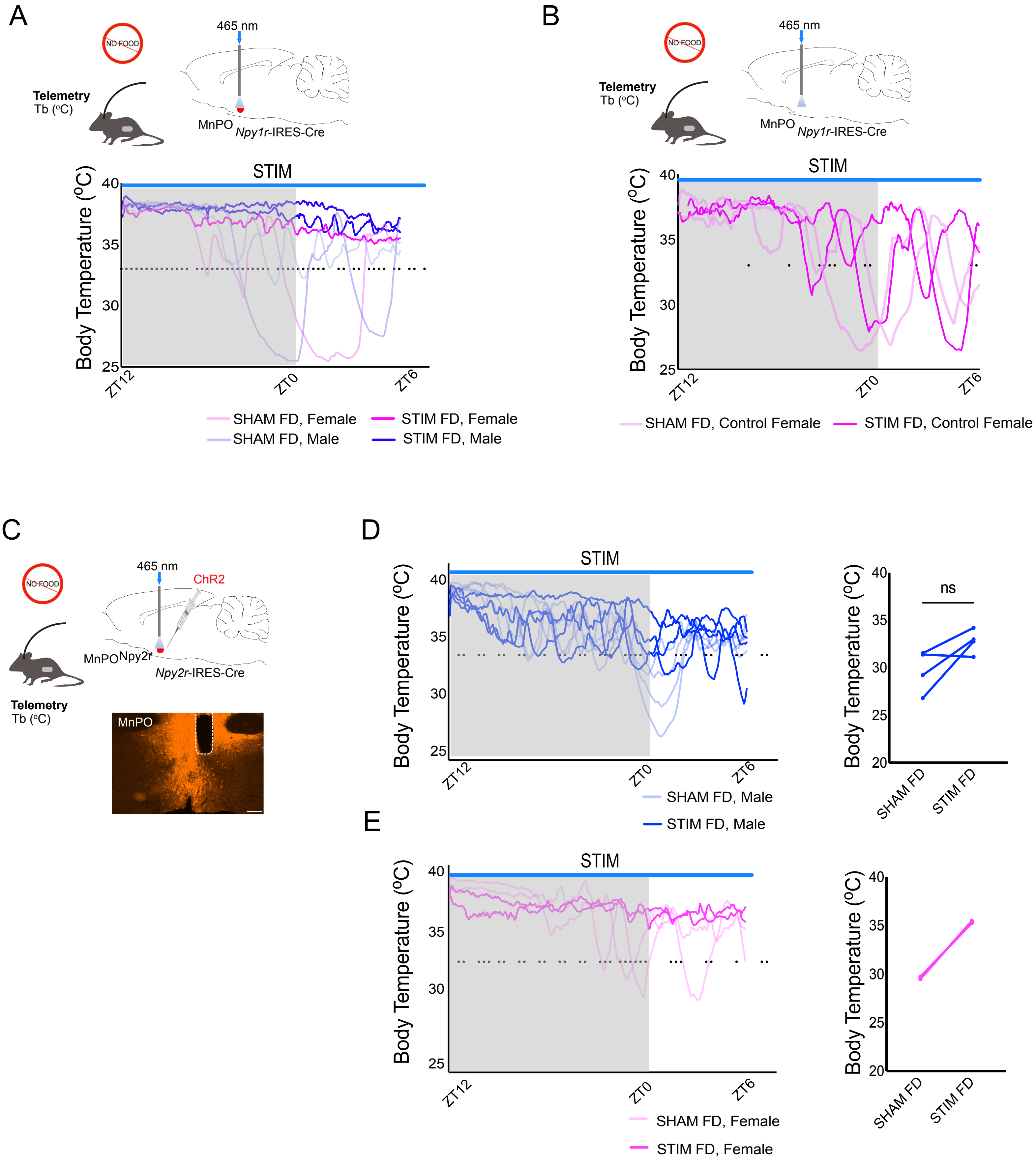

### Ext data Fig 12

#### *A central circuit for energy conservation*

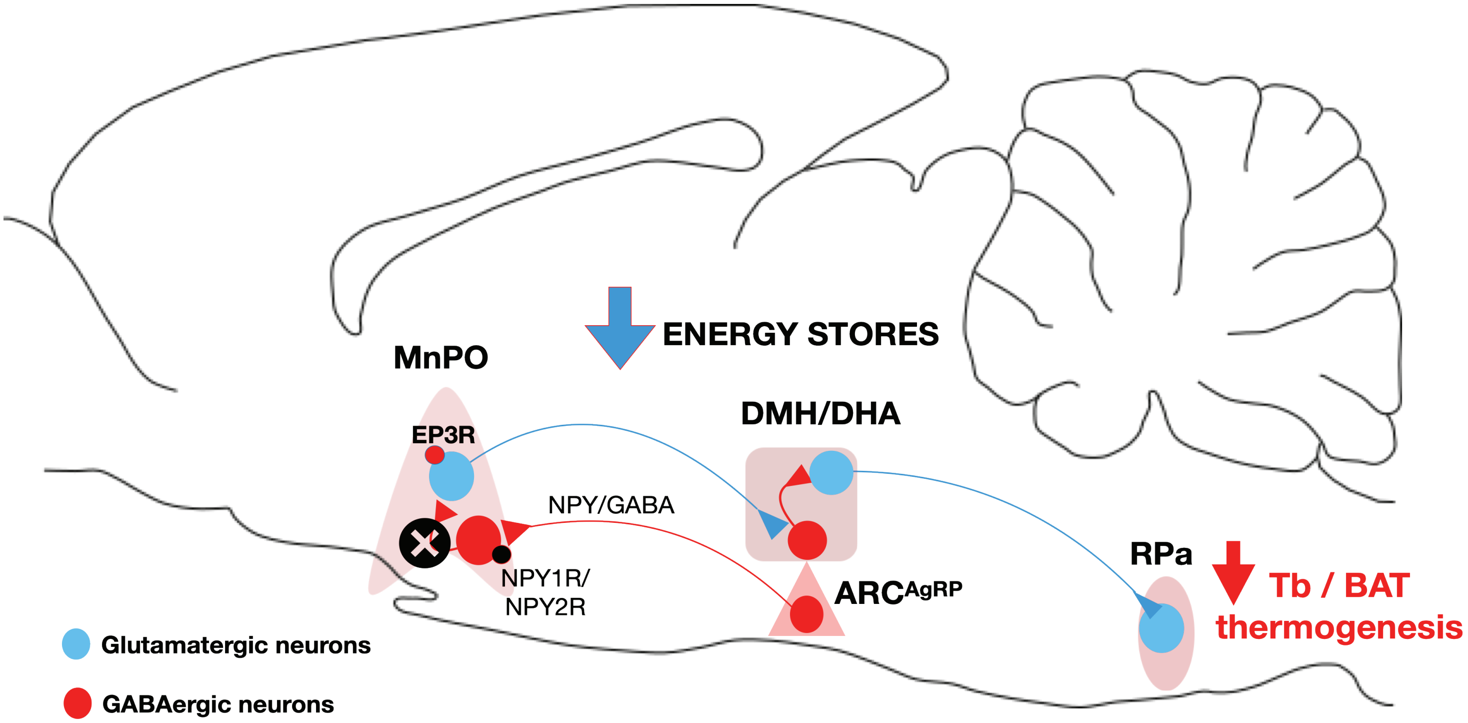

### xExt data Fig 6

#### Extended Data Fig. 6

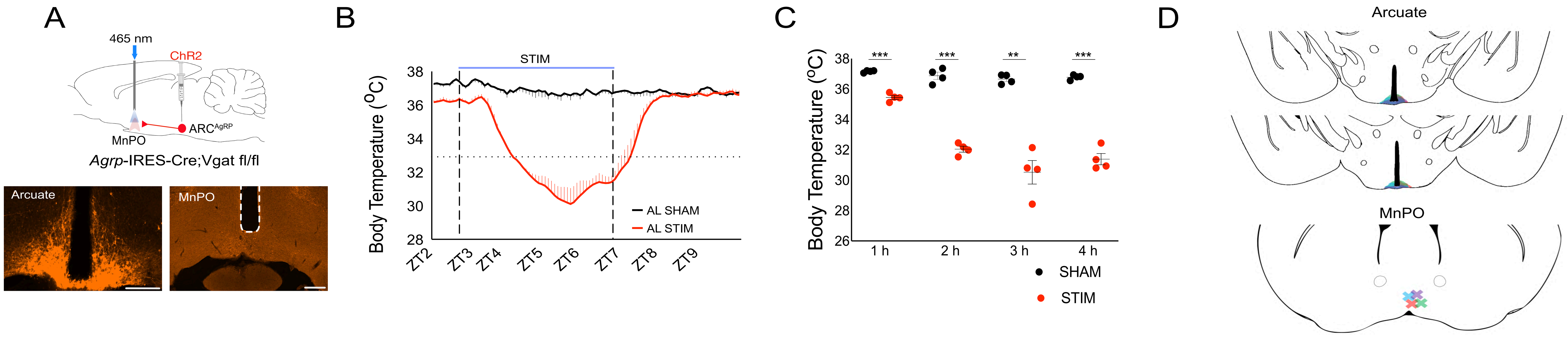
