## Supplementary material for "A Hypothalamic Pathway for Energy Conservation": Ext data Fig 10

A

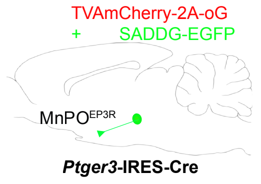


B

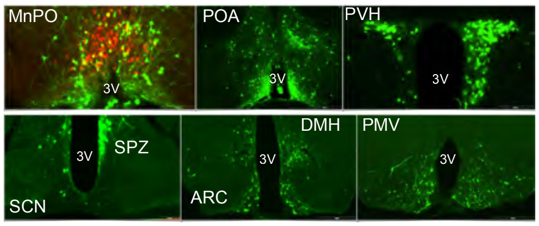


C

| | NM1267 | NM1266 | NM1264 | NM1734 |
|---|---|---|---|---|
| Nucleus accubens | + | None | +++ | + |
| Ventral medial preoptic nucleus (VMPO) | ++ | ++ | ++ | ++ |
| Ventrolateral preoptic nucleus (VLPO) | + | + | + | + |
| Bed nucleus of the stria terminalis (BNST) | ++ | + | ++ | ++ |
| Subfornical organ (SFO) | + | + | + | + |
| Suprachiasmatic nucleus (SCN) | + | + | + | + |
| Subparaventricular zone of the hypothalamus (SPZ) | + | + | + | + |
| Supraoptic nucleus (SON) | + | + | + | None |
| Amygdala | + | + | + | ++ |
| Periventricular hypothalamic nucleus | +++ | +++ | +++ | +++ |
| Paraventricular hypothalamic nucleus | ++ | ++ | +++ | ++ |
| Paraventricular nucleus of the thalamus (PVH) | + | + | + | + |
| Lateroanterior hypothalamic nucleus (LA) | + | + | + | + |
| Reuniens thalamic nucleus | + | + | + | + |
| Lateral hypothalamus (LH) | + | + | + | + |
| Perifornical area (PeF) | + | + | + | + |
| Dorsomedial hypothalamus (dorsal) (DMD) | ++ | + | ++ | ++ |
| Dorsomedial hypothalamus (ventral) (DMV) | +++ | ++ | ++ | + |
| Arcuate nucleus (Arc) | ++ | + | +++ | ++ |
| Tuberomammillary nucleus | + | + | + | + |
| Premammillary nucleus | ++ | ++ | ++ | ++ |
| Dorsal and dorsolateral periaqueductal gray | None | + | + | None |
| Dorsal raphe and median raphe (DR/MnR) | None | + | + | None |
| Pedunculopontine tegmental nucleus (PTg) | + | None | None | None |
| Ventrolateral periaqueductal gray (VLPAG) | + | None | + | + |
| Barrington's nucleus/laterodorsal tegmental nucleus | + | None | None | None |
| Lateral parabrachial nucleus region (LPB) | + | + | None | + |
| Locus coeruleus (LC) | None | None | None | None |
| Raphe pallidus (RPa) | None | None | None | None |
| Parapyrimidal region (PPy) | None | None | None | None |
| Raphe obscurus (Rob) | None | None | None | None |
| Commissural nucleus of the solitary tract/central cana | None | None | None | None |

Sparse innervation + ( < 5 neurons labeled)
Moderate innervation ++ (5-20 neurons labeled)
Dense innervation +++ ( > 20 neurons labeled)

D

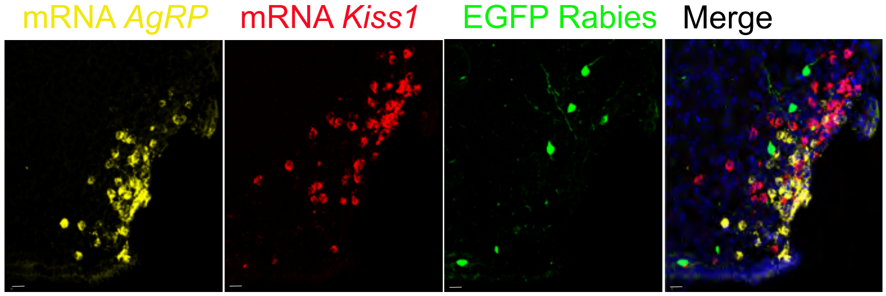
